# Caloric restriction alters lipid metabolism to contribute to tumor growth inhibition

**DOI:** 10.1101/2020.03.09.984302

**Authors:** Evan C. Lien, Anna M. Westermark, Zhaoqi Li, Kiera M. Sapp, Matthew G. Vander Heiden

## Abstract

Dietary interventions can change metabolite levels in the tumor microenvironment, which may then affect cancer cell metabolism to alter tumor growth^1–6^. Although caloric restriction (CR) and the ketogenic diet (KD) are often thought to inhibit tumor growth through lowering blood glucose and insulin levels^7–12^, only CR inhibits the growth of pancreatic ductal adenocarcinoma allografts in mice, demonstrating that this diet can limit tumor growth in other ways. A change in nutrient availability observed with CR, but not the KD, that can contribute to tumor growth inhibition is lower lipid levels in the plasma and in tumor interstitial fluid. Limiting exogenous lipid availability to cultured cancer cells results in up-regulation of stearoyl-CoA desaturase (SCD), an enzyme that converts saturated fatty acids to monounsaturated fatty acids. Fatty acid desaturation is required to dispose of toxic saturated fatty acids, and not because monounsaturated fatty acids are specifically needed for proliferation. Surprisingly, CR also inhibits tumor SCD activity, and enforced SCD expression confers resistance to the effects of CR. Therefore, CR both limits lipid availability and impairs tumor SCD activity, thereby limiting cancer cell adaptation to a diet-induced change in the tumor microenvironment that results in tumor growth inhibition.

---

How diet affects cancer progression and survival is an important question for many cancer patients. While multiple studies have examined how different diets can influence both whole-body hormone signaling and cancer cell signaling processes^7, 8, 10, 11, 13–16^, how dietary interventions might impact tumor growth by altering nutrient access to cancer cells in tumors is less studied. Consistent with the notion that the activities of various metabolic pathways in cancer cells can be strongly influenced by nutrient availability in the local environment^17–21^, dietary deprivation of specific amino acids can limit the growth of some tumors^2, 4–6^. Whether more common dietary modifications that alter the ratios and/or amounts of carbohydrates, proteins, and fats might alter tumor growth via effects on tumor nutrient availability is poorly understood, and one framework for examining the impact of diet on tumor metabolism is to consider how dietary factors modulate the systemic availability of nutrients that are available for use by cancer cells within a tumor^1^. Perturbations to dietary compositions can change metabolite levels in circulation, which in turn can influence metabolite levels in the tumor microenvironment^3^. Thus, diet-induced changes in nutrient availability could affect how cancer cells utilize these nutrients to support cell proliferation, thereby altering tumor growth, progression, and/or response to therapy.

Here, we examine how low carbohydrate diets, such as caloric restriction (CR) and the ketogenic diet (KD), affect tumor metabolism. Many cancer cells and tumors exhibit increased glucose consumption^22, 23^, a phenotype that enables FDG-PET scanning to image cancer in some patients^24^, and low carbohydrate diets are often presumed to inhibit tumor growth in part by lowering blood glucose levels^7–9, 11^. Whether changes in the systemic availability of other nutrients that may alter cancer cell metabolism contribute to the tumor growth inhibitory effects of these diets are much less well studied.

To characterize potential metabolic mechanisms underlying the effects of low carbohydrate diets on tumor growth, we employed a syngeneic mouse cancer model in which AL1376 pancreatic ductal adenocarcinoma (PDAC) cells derived from the *LSL-Kras(G12D)*;*Trp53^fl/fl^*;*Pdx1-Cre* mouse model of pancreatic cancer^25^ are implanted into C57BL/6J mice to form allograft tumors^26^. Since orthotopic implantation of PDAC cells into the pancreas leads to pancreatic exocrine insufficiency and impaired food digestion^26^, we compared subcutaneous tumor growth in mice exposed to different diets. Specifically, after the formation of palpable tumors, mice were switched to either a control diet, CR, or a KD. The CR regimen examined involves a 40% reduction in caloric consumption through limitation of the carbohydrate portion of the diet, whereas the KD is 90% fat, 9% protein, and 1% carbohydrates (see Extended Data Table 1 for composition of all diets used in this study). Interestingly, only CR, but not the KD, inhibits tumor growth in this model, as measured by tumor volume over time and tumor weights at study endpoint (Fig. 1a-d, Extended Data Fig. 1a-d). Both CR and the KD result in reduced animal body weights (Extended Data Fig. 1e-f), but the tumor growth inhibitory effects of CR cannot be explained by a loss of body weight resulting from a reduction in calories consumed since tumor growth is still inhibited by CR even when normalized to the animals’ body weights (Fig. 1a-d). Additionally, we confirmed that mice fed the KD are not calorically restricted. Although they consume less food daily by weight (Extended Data Fig. 1g), they consume similar amounts of calories compared to control when accounting for the higher caloric density of the KD (Extended Data Fig. 1h, Extended Data Table 1). Despite these divergent effects on tumor growth, both CR and the KD result in a similar decrease in blood glucose levels (Fig. 1e-f). Moreover, only the KD significantly reduces fasting plasma insulin levels, although CR results in a trend toward decreased fasting plasma insulin that is not statistically significant (Fig. 1g-h). Taken together, these results indicate that different low carbohydrate diets cannot be assumed to be equivalent in their effects on tumor growth, even if they decrease blood glucose and insulin.

**Figure 1.**
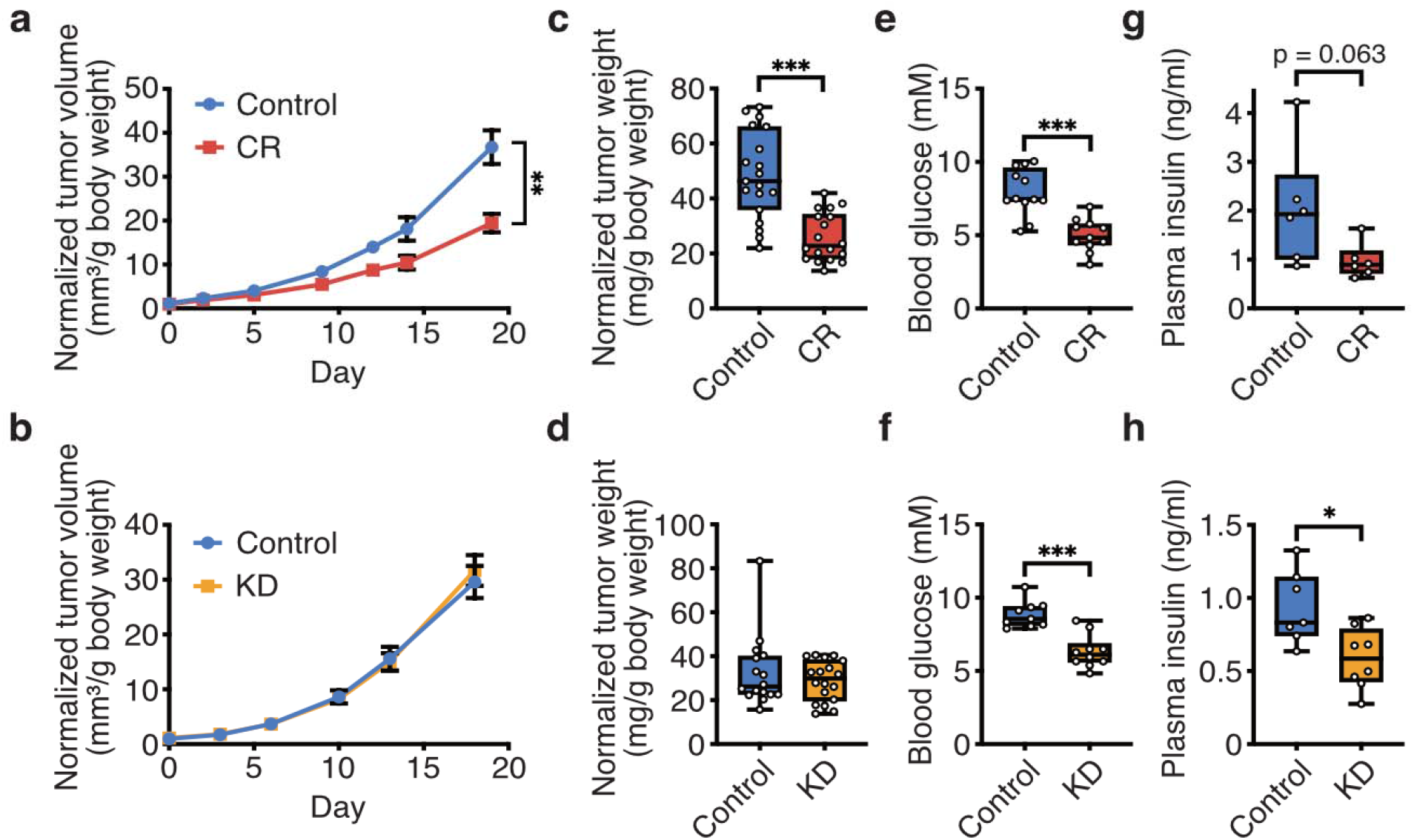
Caloric restriction, but not the ketogenic diet, impairs growth of pancreatic cancer allograft tumors. **a**, **b**, Normalized tumor volumes of subcutaneous AL1376 pancreatic ductal adenocarcinoma allografts implanted in C57BL/6J mice exposed to (**a**) CR or a control diet, or (**b**) the KD or a control diet. Tumor volumes normalized to animal body weight are shown. Data presented are mean ± SEM; (**a**) Control n = 4 male + 8 female mice, CR n = 3 male + 8 female mice; (**b**) Control n = 5 male + 4 female mice, KD n = 5 male + 5 female mice. Two-way repeated measures analysis of variance (ANOVA) was used for comparison between groups. **c**, **d**, Tumor weights normalized to animal body weight of subcutaneous AL1376 allografts harvested at endpoint for the experiment shown in (**a-b**) where mice were exposed to (**c**) CR or a control diet, or (**d**) the KD or a control diet. Data are presented as box-and-whisker plots displaying median and interquartile ranges; (**c**) Control n = 4 male + 8 female mice, CR n = 3 male + 8 female mice; (**d**) Control n = 5 male + 4 female mice, KD n = 5 male + 5 female mice. A two-tailed Student’s t-test was used for comparison between groups. **e-h**, Effects of control diets, CR, or the KD on (**e-f**) blood glucose levels or (**g-h**) fasting plasma insulin levels as indicated. Data are presented as box-and-whisker plots displaying median and interquartile ranges; (**e**) Control n = 12, CR n = 11; (**f**) Control n = 9, KD n = 10; (**g**) Control n = 6, CR n = 6; (**h**) Control n = 7, KD n = 8. A two-tailed Student’s t-test was used for comparison between groups. *P < 0.05, **P < 0.01, ***P < 0.001.

The finding that CR inhibits tumor growth while having a similar effect as the KD on blood glucose levels, and has a less pronounced effect on fasting plasma insulin levels, suggests that the growth inhibitory effects of CR may not be fully explained by its effects on blood glucose and insulin. Since dietary perturbations may induce broad changes in the systemic availability of metabolites beyond those altered in the diet itself^1^, we reasoned that these two diets may have distinct effects on the systemic availability of other metabolites, and that a change in the availability of other nutrient(s) might contribute to the tumor growth inhibition observed in mice exposed to CR.

To determine how nutrient availability in tumors is differentially affected by each diet, we measured the levels of several metabolites in the plasma and tumor interstitial fluid (TIF) harvested from tumor-bearing mice fed a control diet, the CR diet, or the KD. TIF comprises of the fluid surrounding cancer cells within the tumor microenvironment, and therefore may better reflect the nutrient environment to which cancer cells are exposed in tumors^3^. For example, despite lowering blood glucose levels, the KD does not lower glucose levels in TIF (∼2.8 mM for both control and KD), whereas a decrease in TIF glucose levels from ∼1.6 mM to ∼0.8 mM is observed when mice are exposed to CR (Fig. 2a-b). When cultured at these glucose concentrations *in vitro*, the proliferation of the PDAC cells is minimally impaired at the average CR TIF concentration of 0.8 mM, although this difference is not statistically significant (Fig. 2c). Therefore, even though both CR and the KD lower blood glucose levels, CR may be more effective at reducing glucose availability to cells within tumors, which may contribute in part to its growth inhibitory effects. However, since any effect on cell proliferation at 0.8 mM glucose is small, it remains possible that changes in the availability of other nutrients contribute to the anti-tumor growth effects of CR.

**Figure 2.**
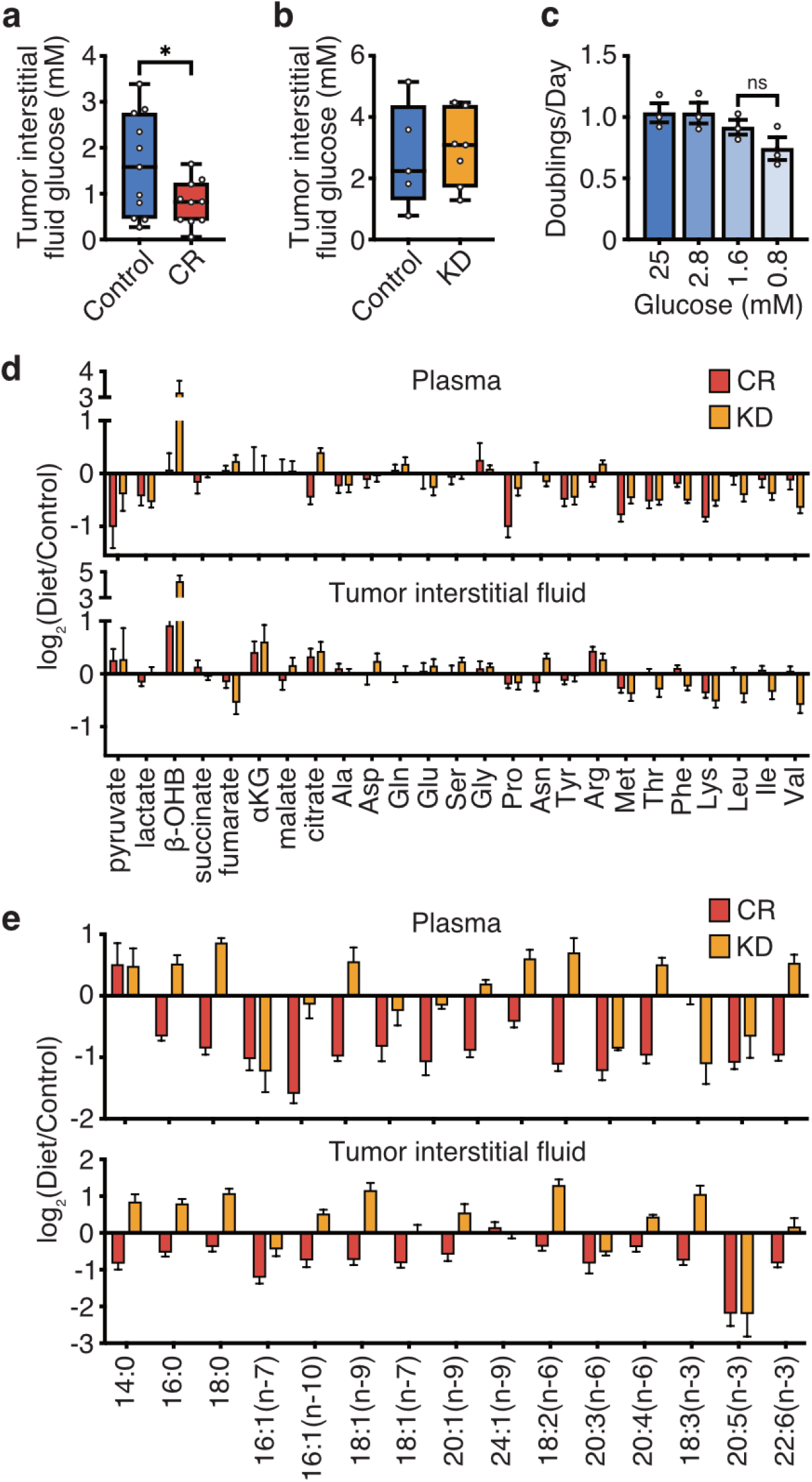
Caloric restriction and the ketogenic diet differentially alter nutrient levels in the plasma and tumor interstitial fluid. **a**, **b**, Concentration of glucose in tumor interstitial fluid collected from subcutaneous AL1376 allograft tumors in C57BL/6J mice exposed to (**a**) CR or a control diet, or (**b**) the KD or a control diet. Data are presented as box-and-whisker plots displaying median and interquartile ranges; (**a**) Control n = 11, CR n = 10; (**b**) Control n = 5, KD n = 7. A two-tailed Student’s t-test was used for comparison between groups. **c**, Doubling times of AL1376 cells cultured in media containing the indicated concentrations of glucose. Data are presented as mean ± SEM; n = 3 biologically independent replicates per group. A two-tailed Student’s t-test was used for comparison between groups. **d**, Fold change in the specified metabolite levels induced by CR or the KD relative to the control diet measured in the plasma (upper) or tumor interstitial fluid (lower) as indicated. Data are presented as mean ± SEM; Plasma: CR n = 8, KD n = 9; Tumor interstitial fluid: CR n = 13, KD n = 6. **e**, Fold changes in the specified fatty acid levels induced by CR or the KD relative to the control diet measured in the plasma (upper) or tumor interstitial fluid (lower). Data are presented as mean ± SEM; Plasma: CR n = 7, KD n = 4; Tumor interstitial fluid: CR n = 10, KD n = 4. *P < 0.05, **P < 0.01, ***P < 0.001.

Levels of most metabolites measured that are involved in central carbon and amino acid metabolism are not robustly changed in the plasma or TIF by either CR or the KD relative to the control diet (Fig. 2d). Consistent with previous measurements of serum amino acid changes induced by the KD^27^, levels of essential amino acids are consistently reduced by the KD relative to the control diet in both plasma and TIF (Fig. 2d), likely due to the lower protein content of the KD relative to the control diet (Extended Data Table 1). CR also affects levels of some essential amino acids, but both the number of affected amino acids and the magnitude of change are less than that observed in the KD and are more prominent in plasma than in TIF. Another major difference between CR and the KD is that the KD is more effective at raising the levels of the β -hydroxybutyrate (β -OHB). Of note, some cancer cells can consume and metabolize ketones, and knockdown of ketone metabolism genes can sensitize tumors to a KD^28^. Therefore, the ability of the KD to robustly increase ketone levels in the plasma and TIF may contribute to why this diet does not inhibit tumor growth in this model.

To measure the availability of fatty acids in the plasma and TIF, lipids were derivatized into fatty acid methyl esters and measured by gas chromatography-mass spectrometry (GC-MS) (Fig. 2e). A striking difference in fatty acids was observed in both plasma and TIF when mice were exposed to CR versus the KD. Most fatty acids measured are increased by the KD relative to the control diet, consistent with the KD consisting of 90% fat (Extended Data Table 1). In contrast, levels of almost all fatty acids are reduced by CR, particularly in the TIF, which is consistent with previous studies showing how CR can decrease circulating lipid levels^29–33^. This observation suggests that compared to the KD, decreased systemic availability of lipids and fatty acids to the tumor is specific to CR, and environmental lipid limitation could be one metabolic mechanism by which CR inhibits tumor growth.

Since changes to the nutrient environment can alter the metabolism of cancer cells, we first asked how lipid deprivation impacts cellular fatty acid metabolism *in vitro*. When cultured in lipid-depleted media in which only trace amounts of fatty acids remain^34^ (Extended Data Fig. 2a), AL1376 cells retain the ability to proliferate, albeit at a slower rate (Fig. 3a). A variety of cancer cells derived from multiple tissues can also proliferate in lipid-depleted media, including: LGSP (a lung adenocarcinoma cell line derived from the *LSL-Kras(G12D)*;*Trp53^fl/fl^*;*Ad-Cre* mouse lung cancer model^35, 36^), HeLa (human cervical cancer), Panc1 (human PDAC), and A549 (human lung adenocarcinoma) cells (Extended Data Fig. 2b). To begin to characterize the metabolic consequences of culture in these conditions, we measured the fatty acid composition of lipid-starved cells. Under these conditions, all cell lines tested have decreased levels of polyunsaturated fatty acids (PUFAs), which is expected since these are essential fatty acids that cannot be synthesized by mammalian cells *de novo* and must be obtained from the environment (Fig. 3b, Extended Data Fig. 2c). In contrast, levels of saturated fatty acids (SFAs) and monounsaturated fatty acids (MUFAs), which can be synthesized *de novo*, are relatively maintained, with only slight reductions in some species (Fig. 3b, Extended Data Fig. 2c). These data are consistent with exogenous lipid limitation promoting *de novo* fatty acid synthesis to support continued proliferation, and indeed lipid-starved cells have an increased rate of fatty acid synthesis as measured by ^13^C-glucose and ^13^C-glutamine incorporation into cellular fatty acids followed by rate calculations using Fatty Acid Source Analysis^37^ (Fig. 3c, Extended Data Fig. 2d, Extended Data Fig. 3, Supplementary Table 1).

**Figure 3.**
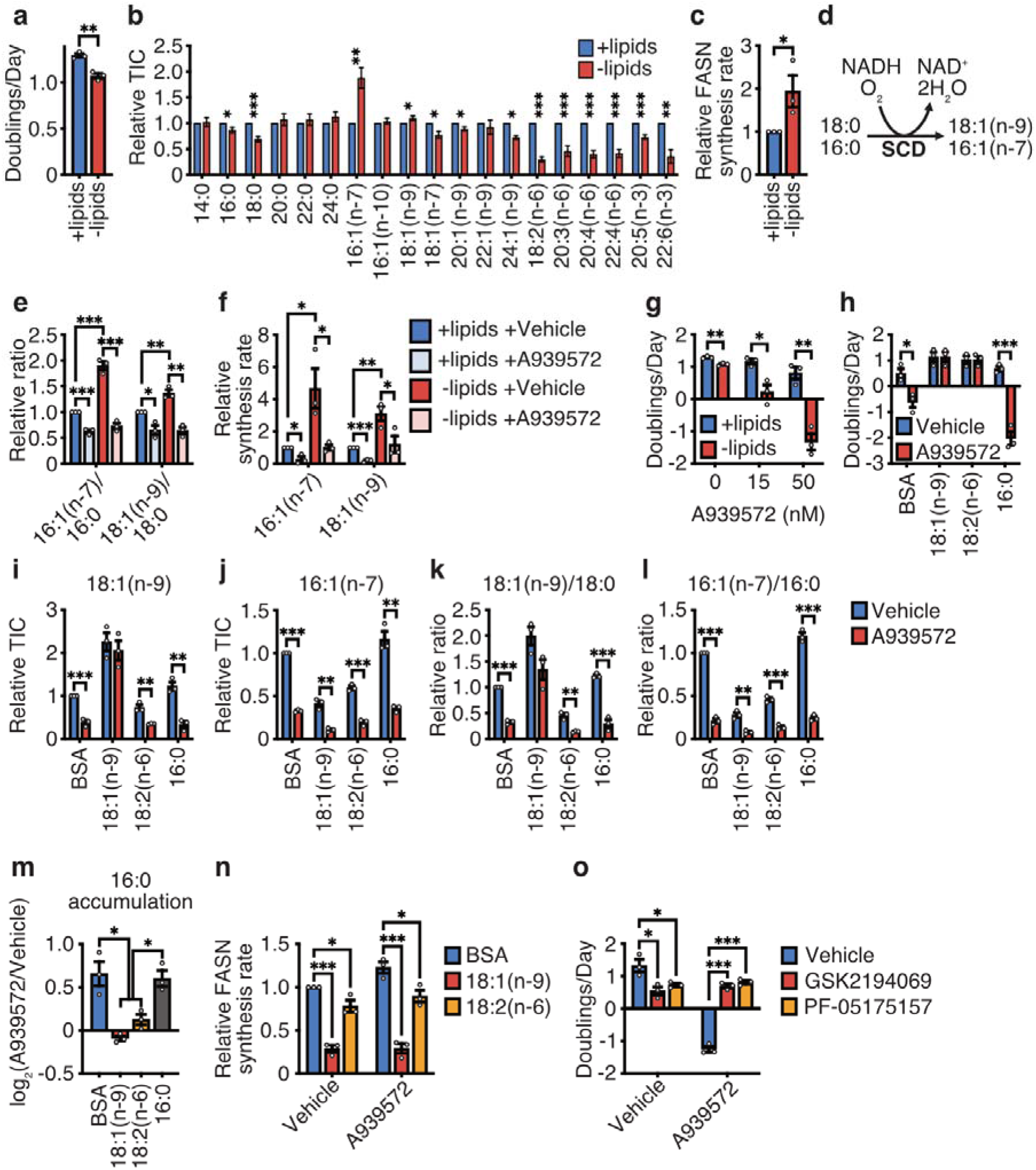
Increased stearoyl-CoA desaturase activity is required for cancer cells to adapt to exogenous lipid limitation. **a**, Doubling times of AL1376 cells cultured in lipidated versus de-lipidated media. **b**, Fatty acid levels in AL1376 cells cultured in lipidated versus de-lipidated media for 24 h. n = 5 biologically independent replicates. **c**, AL1376 cells were cultured in the presence of U-^13^C-glucose and U-^13^C-glutamine for 24 h in lipidated versus de-lipidated media, and ^13^C incorporation into fatty acids was detected by GC-MS. The mass isotopologue distributions and total pool sizes of detected fatty acids were used to calculate a FASN synthesis rate by fatty acid source analysis (FASA, see Supplementary Table 1). A one-tailed Student’s t-test was used for comparison between groups. **d**, Schematic of the reaction catalyzed by stearoyl-CoA desaturase (SCD). **e**, **f**, MUFA/SFA ratios (**e**) and synthesis rates of 16:1(n-7) and 18:1(n-9) (**f**), calculated as described in (**c**) (see also Supplementary Table 1), measured in AL1376 cells cultured in lipidated versus de-lipidated media, with or without 50 nM of the SCD inhibitor A939572 for 24 h as indicated. **g**, Doubling times of AL1376 cells cultured in lipidated versus de-lipidated media with the indicated concentrations of A939572. **h**, Doubling times of AL1376 cells cultured in de-lipidated media with or without 50 nM A939572, BSA, 50 µM 18:1(n-9), 50 µM 18:2(n-6), or 15 µM 16:0 as indicated. **i-l**, 18:1(n-9) (**i**), 16:1(n-7) (**j**), 18:1(n-9)/18:0 (**k**), and 16:1(n-7)/16:0 (**l**) levels measured in AL1376 cells cultured in de-lipidated media with or without 50 nM A939572, BSA, 50 µM 18:1(n-9), 50 µM 18:2(n-6), or 15 µM 16:0 for 48 h as indicated. **m**, Fold change in levels of 16:0 induced by treatment with 50 nM A939572 in AL1376 cells cultured in de-lipidated media containing BSA, 50 µM 18:1(n-9), 50 µM 18:2(n-6), or 15 µM 16:0 for 48 h as indicated. **n**, FASN synthesis rates, determined as described in (**c**) (see also Supplementary Table 1), measured in AL1376 cells cultured in de-lipidated media with or without 50 nM A939572, BSA, 50 µM 18:1(n-9), or 50 µM 18:2(n-6) as indicated. **o**, Doubling times of AL1376 cells cultured in de-lipidated media containing 50 nM A939572, 0.3 µM of the FASN inhibitor GSK2194069, or 10 µM of the ACC inhibitor PF-05175157. All data are presented as mean ± SEM; unless otherwise indicated, n = 3 biologically independent replicates, and a two-tailed Student’s t-test was used for comparison between groups. *P < 0.05, **P < 0.01, ***P < 0.001.

Although lipid limitation either has minimal effect on or decreases the levels of most fatty acid species, levels of the 16:1(n-7) MUFA are markedly increased in cells cultured in lipid-depleted conditions (Fig. 3b, Extended Data Fig. 2c). The enzyme stearoyl-CoA desaturase (SCD) is responsible for the synthesis of the 16:1(n-7) and 18:1(n-9) MUFAs from the 16:0 and 18:0 SFAs, respectively (Fig. 3d). The 16:1(n-7)/16:0 and 18:1(n-9)/18:0 ratios, which have been used as surrogates of SCD activity^38, 39^, are also significantly increased in lipid-starved cells, and these ratios are decreased by the SCD inhibitor A939572 (Fig. 3e, Extended Data Fig. 2e-f). Direct measurement of ^13^C-glucose and ^13^C-glutamine incorporation into 16:1(n-7) and 18:1(n-9) also suggests increased flux through SCD in lipid-starved cells, which is inhibited by A939572 (Fig. 3f, Extended Data Fig. 2g-h, Extended Data Fig. 3c-d, Supplementary Table 1). Together, these data suggest that SCD activity is increased when exogenous lipids are limited.

To determine whether increased SCD activity is necessary for cells to respond to lipid starvation, we evaluated how SCD inhibition affects cell proliferation. Whereas cells are minimally affected by A939572 in lipid-replete media, in lipid-depleted media SCD inhibition strongly impairs cell proliferation in all cell lines and even induces cell death in AL1376, LGSP, and HeLa cells (Fig. 3g, Extended Data Fig. 2i). These data argue that up-regulation of SCD activity is necessary for cancer cells when exogenous lipids are less abundant.

A recent report demonstrated that fatty acid desaturation can support the regeneration of the co-factor NAD^+^, which is a product of these reactions (Fig. 3d), although this study focused specifically on fatty acid desaturase 1 (FADS1) and fatty acid desaturase 2 (FADS2)^40^. Cancer cells require NAD^+^ for proliferation because it is necessary for the oxidative synthesis of metabolites such as aspartate, serine, and nucleotides^41–44^. We therefore considered the possibility that lipid-limited cells may up-regulate SCD activity to regenerate sufficient NAD^+^ to support proliferation. Perturbations that inhibit proliferation by decreasing the NAD^+^/NADH ratio can be rescued by the addition of exogenous pyruvate, which regenerates NAD^+^ through its conversion to lactate^41, 45^. Therefore, we assessed whether pyruvate could rescue the effects of A939572 on cell growth and viability (Extended Data Fig. 4a-e). In all cells tested, pyruvate does not rescue SCD inhibition, suggesting that NAD^+^ limitation is not the mechanism by which SCD inhibition impairs cell growth and viability.

SCD is thought to be important to produce MUFAs for cancer cell proliferation^46–49^. Cancer cells have variable sensitivities to SCD inhibition, and some resistant cells can make an alternative MUFA, sapienate (16:1(n-10)), that is synthesized by FADS2 to maintain MUFA levels in cells^50^. However, despite variable sensitivity to SCD inhibition among the cells considered here, in these cells the levels of 16:1(n-10) were minimally increased upon A939572 treatment (Extended Data Fig. 5a), and no change was observed in the 16:1(n-10)/16:0 ratio or synthesis rate of 16:1(n-10) (Extended Data Fig. 5b-c, Supplementary Table 1). Thus, the lack of MUFAs could be responsible for the effects of SCD inhibition on the proliferation and viability of these cells.

Consistent with prior studies^46, 48, 49, 51^, exogenous supplementation with the 18:1(n-9) MUFA rescues cell death induced by SCD inhibition in both AL1376 and LGSP cells (Fig. 3h, Extended Data Fig. 6a). Exogenous 18:1(n-9) in the presence of A939572 rescues levels of 18:1(n-9) but not 16:1(n-7) (Fig. 3i-j, Extended Data Fig. 6c-d) and also rescues the 18:1(n-9)/18:0 ratio, but not 16:1(n-7)/16:0 ratio (Fig. 3k-l, Extended Data Fig. 6e-f). Similar observations in other studies have led to the conclusion that SCD inhibition impairs cell proliferation and viability by depleting MUFA levels and/or by lowering the MUFA/SFA ratios within cells^46, 48, 49^. However, we noted that SCD not only synthesizes MUFAs, but also consumes the SFAs 16:0 and 18:0 which are toxic to cells^52–54^ (Fig. 3d). The increase in 16:1(n-7) levels in lipid-starved cells (Fig. 3b, Extended Data Fig. 2c) raises the possibility that under conditions of exogenous lipid starvation, an important function of SCD may be to desaturate SFAs, in particular newly synthesized 16:0 into 16:1(n-7). In support of this hypothesis, SCD inhibition with A939572 leads to the accumulation of the 16:0 SFA, and this accumulation is reversed with the supplementation of exogenous 18:1(n-9) (Fig. 3m, Extended Data Fig. 6g).

To determine whether SCD inhibition is toxic to cells because it depletes MUFAs and/or lowers the MUFA/SFA ratio, or if SCD activity is required to limit accumulation of toxic SFAs such as 16:0, we supplemented cells with 18:2(n-6), an essential PUFA that mammalian cells cannot synthesize *de novo*. Of note, providing 18:2(n-6) rescues the toxicity of SCD inhibition with A939572 in AL1376 and LGSP cells (Fig. 3h, Extended Data Fig. 6a). We confirmed that 18:2(n-6) also rescues the effects of SCD inhibition in HeLa, Panc1, and A549 cells (Extended Data Fig. 6b). Since cells cannot convert 18:2(n-6) into MUFAs, 18:2(n-6) supplementation does not rescue levels of 18:1(n-9) or 16:1(n-7), or the 18:1(n-9)/18:0 or 16:1(n-7)/16:0 ratios in cells treated with A939572 (Fig. 3i-l, Extended Data Fig. 6c-f). However, 18:2(n-6) does block the accumulation of 16:0 that is induced by SCD inhibition (Fig. 3m, Extended Data Fig. 6g). Therefore, these data are most consistent with the acute toxicity of SCD inhibition being mediated by 16:0 accumulation, rather than a reduction of intracellular MUFAs or the decrease in the MUFA/SFA ratios. To corroborate this, supplementation with exogenous 16:0 in combination with A939572 exacerbates the toxicity of SCD inhibition (Fig. 3h, Extended Data Fig. 6a). 16:0 has no effect on MUFA levels or the MUFA/SFA ratios, while still leading to increased levels of 16:0 (Fig. 3i-m, Extended Data Fig. 6c-g).

We next asked how 18:1(n-9) and 18:2(n-6) prevent the accumulation of 16:0 following SCD inhibition. Since 16:0 is the first fatty acid synthesized by the *de novo* fatty acid synthesis pathway, we reasoned that the presence of these exogenous fatty acids may inhibit fatty acid production through FASN. Previous studies have shown that in a lipid-replete environment, the activity of the SREBP transcription factors is repressed, which leads to the down-regulation of fatty acid synthesis genes^55^. Indeed, by tracing ^13^C-glucose and ^13^C-glutamine into fatty acids, we find that both 18:1(n-9) and 18:2(n-6) inhibit fatty acid synthesis rates when cells are treated with either vehicle or A939572 (Fig. 3n, Extended Data Fig. 6h, Supplementary Table 1). These data suggest 18:1(n-9) and 18:2(n-6) prevent 16:0 accumulation upon SCD inhibition by inhibiting the *de novo* synthesis of 16:0 through FASN.

To further test whether toxicity from SCD inhibition is mediated by the accumulation of SFAs, we determined whether orthogonal methods of inhibiting fatty acid synthesis would rescue the toxicity of SCD inhibition. Of note, a recent screen identified the loss of acetyl-CoA carboxylase 1 (ACACA), which is required for *de novo* fatty acid synthesis, to be protective against 16:0 toxicity^54^. We utilized GSK2194069, a fatty acid synthase inhibitor^56^, and PF-05175157, an acetyl-CoA carboxylase (ACC) inhibitor^57^, at concentrations that only slightly impair cell proliferation in lipid-depleted media, either alone or in combination with the SCD inhibitor A939572. Of note, while each inhibitor either impairs cell proliferation or induces cell death on its own, inhibition of FASN or ACC rescues the toxicity of SCD inhibition (Fig. 3o). Similar results are observed in LGSP, HeLa, Panc1, and A549 cells (Extended Data Fig. 6i). That *de novo* fatty acid synthesis inhibitors antagonize SCD inhibition highlights the importance of understanding how distinct lipid metabolism pathways interact with each other, especially given the ongoing interest in developing cancer therapies that target various nodes in lipid metabolism^58–63^. Taken together, these data confirm that an important reason for the acute toxicity of SCD inhibition under lipid-starved conditions is the accumulation of 16:0 SFA, rather than the depletion of MUFAs or alterations in the MUFA/SFA ratios.

Modeling exogenous lipid limitation *in vitro* revealed that the up-regulation of SCD activity is a critical adaptation for lipid-starved cancer cells. To determine whether this finding might provide insight into how CR limits tumor growth, we assessed the activity of SCD in AL1376 tumors harvested from calorically restricted mice. Of note, CR decreases the levels of 16:1(n-7) and 18:1(n-9) MUFAs (Fig. 4a) and the 16:1(n-7)/16:0 and 18:1(n-9)/18:0 ratios (Fig. 4b). These data suggest that CR impairs SCD activity in tumors, and consistently we found that calorically restricted tumors express lower levels of the SCD protein (Fig. 4c), despite the fact that CR results in lipid depletion in the tumor microenvironment (Fig. 2e).

**Figure 4.**
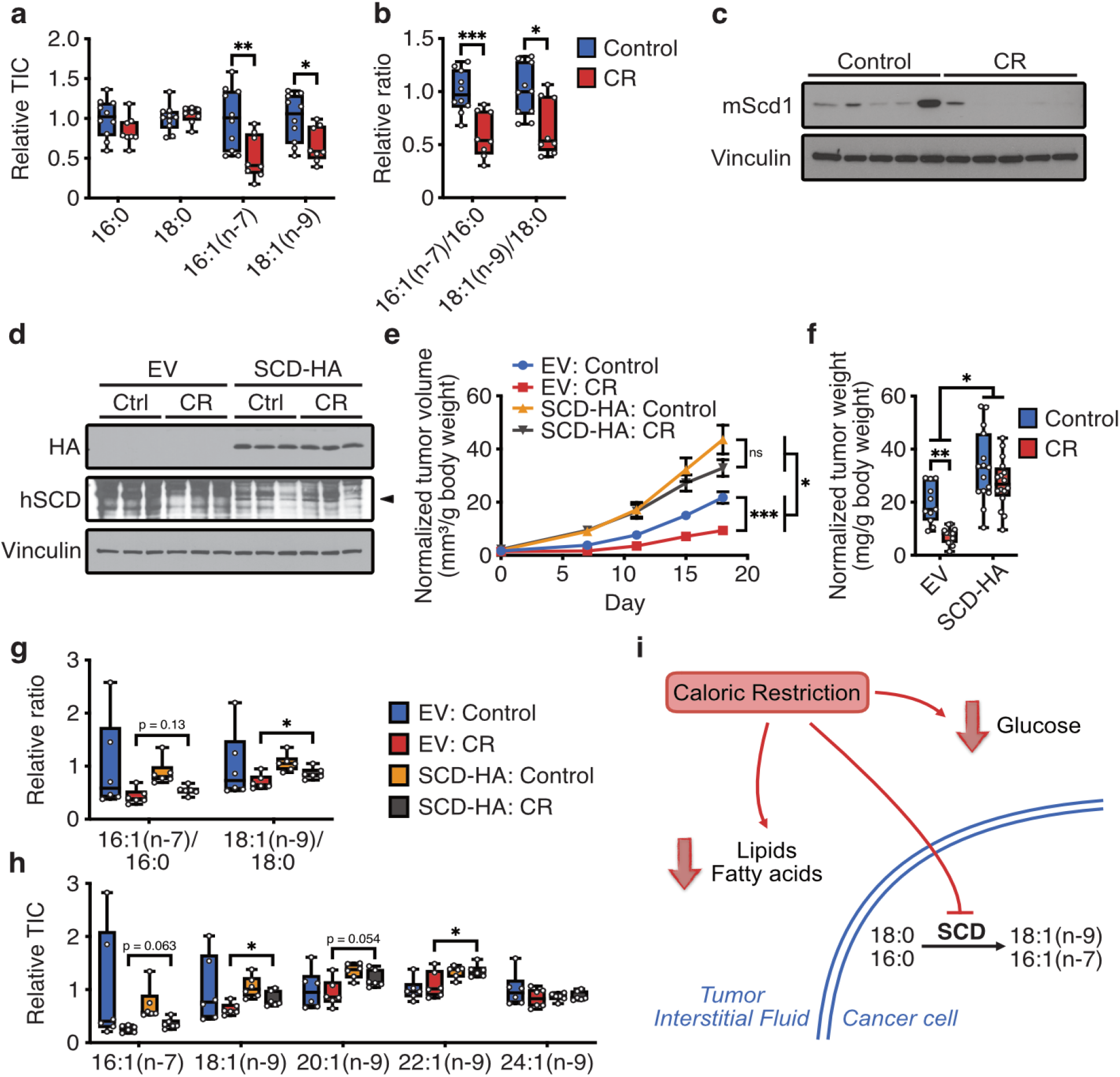
Caloric restriction inhibits tumor growth in part by limiting exogenous lipid availability and impairing tumor SCD activity. **a**, **b**, The indicated fatty acid levels (**a**) and MUFA/SFA ratios (**b**) measured in subcutaneous AL1376 allograft tumors harvested from C57BL/6J mice exposed to a control or CR diet. Data are presented as box-and-whisker plots displaying median and interquartile ranges; Control n = 10, CR n = 8. A two-tailed Student’s t-test was used for comparison between groups. **c**, Immunoblot for mouse Scd1 (mScd1) and vinculin in lysates from subcutaneous AL1376 allograft tumors harvested from C57BL/6J mice exposed to a control or CR diet. **d**, Immunoblot for HA, human SCD (hSCD), and vinculin in lysates from subcutaneous AL1376 allograft tumors expressing an empty vector (EV) or SCD-HA harvested from C57BL/6J mice exposed to a control or CR diet. **e**, **f**, Tumor volumes (**e**) and tumor weights (**f**) normalized to animal body weight of subcutaneous AL1376 allograft tumors expressing EV or SCD-HA in C57BL/6J mice exposed to a control or CR diet for 18 days. Data are presented as (**e**) mean ± SEM and (**f**) box-and-whisker plots displaying median and interquartile ranges; EV: Control n = 3 male + 3 female mice, EV: CR n = 4 male + 4 female mice, SCD-HA: Control n = 4 male + 4 female mice, SCD-HA: CR n = 4 male + 4 female mice. Two-way repeated measures analysis of variance (ANOVA) was used for comparison between groups. **g**, **h**, MUFA/SFA ratios (**g**) and MUFA levels (**h**) measured in subcutaneous AL1376 allograft tumors expressing EV or SCD-HA from C57BL/J6 mice exposed to a control or CR diet for 7 days. Data are presented as box-and-whisker plots displaying median and interquartile ranges; n = 6 per group. A two-tailed Student’s t-test was used for comparison between groups. **i**, Model depicting the effects of caloric restriction on the exogenous availability of lipids, fatty acids, and glucose, as well as cancer cell SCD activity, as contributors to how this diet can impact tumor growth. *P < 0.05, **P < 0.01, ***P < 0.001.

Together, these observations suggest a model in which CR limits lipid availability to the tumor and also impairs SCD activity, therefore preventing cancer cell adaptation to a depleted lipid environment (Fig. 4i). To test whether this combination of lipid limitation and SCD inhibition contributes to how CR inhibits tumor growth, we subcutaneously implanted AL1376 cells engineered to express SCD into mice that were administered either a control or CR diet. Exogenous SCD expression in tumors was confirmed by Western blot (Fig. 4d). Interestingly, even under the control diet, tumors with increased SCD expression grow faster relative to control, suggesting that lipid availability may be limiting for tumor growth even when mice are fed a control diet (Fig. 4e-f, Extended Data Fig. 7a-b). Most importantly, SCD-expressing tumors grow faster relative to control in calorically restricted mice, indicating that SCD activity can promote the growth of calorically restricted tumors (Fig. 4e-f, Extended Data Fig. 7a-b). Moreover, SCD expression partially rescues the tumor growth inhibitory effects of CR. When comparing the raw tumor volumes and weights, CR inhibits the growth of control tumors by ∼70%, whereas the growth of SCD-expressing tumors is inhibited by ∼40% (Extended Data Fig. 7a-b). This ∼40% inhibition is accounted for by the 40% reduction in calories in the CR feeding regimen, and when tumor volumes and weights are normalized to animal body weights, the inhibitory effects of CR are almost completely rescued (Fig. 4e-f). These data argue that the inhibition of tumor growth by CR that extends beyond that accounted for by the reduction in caloric intake is partially rescued by the enforced expression of SCD. This observation is consistent with CR impairing tumor growth, at least in part, by both decreasing lipid availability and inhibiting tumor SCD activity (Fig. 4i).

Finally, to confirm increased SCD activity in SCD-expressing tumors, we measured MUFA/SFA ratios and the levels of MUFAs in these tumors. Interestingly, in tumors collected at the endpoint of the experiment 18 days after diet initiation, CR is equally effective at lowering 16:1(n-7)/16:0, 18:1(n-9)/18:0, 16:1(n-7) and 18:1(n-9) levels for both control and SCD-expressing tumors (Extended Data Fig. 7c-d). However, several factors may be masking the effects of SCD expression in the cancer cells within the tumors. First, there is a high degree of cellular heterogeneity in PDAC tumors, even when grown subcutaneously^26^, and our fatty acid measurements are conducted on bulk tumors containing cancer cells, stromal cells, extracellular matrix, and tumor interstitial fluid. Because the SCD-expressing cancer cells may constitute only a percentage of this bulk population, cancer cell increases in MUFA/SFA and MUFA levels may be masked. Second, endpoint tumors are large with variable necrotic regions, and hypoxia, which can affect the oxygen-requiring SCD reaction^64, 65^, would be expected to affect MUFA production in poorly vascularized regions of the tumor. In an attempt to overcome these issues, we evaluated MUFA/SFA ratios and the levels of MUFAs in tumors collected from mice 7 days after diet initiation, a time point at which tumors are smaller and do not contain necrotic regions. Here, the MUFA/SFA ratios are partially rescued in the SCD-expressing versus control tumors from calorically restricted mice (Fig. 4g). Moreover, SCD-expressing tumors also have higher levels of MUFAs (Fig. 4h). Together, this suggests that the SCD-expressing tumors, which are partially resistant to the growth inhibitory effects of CR, have evidence of increased SCD activity.

By exploring how diet modulates nutrient availability to impact tumor metabolism and progression, we have uncovered a metabolic mechanism by which CR can inhibit tumor growth. Not only does CR lower circulating glucose levels, but it can also decrease lipid availability and SCD activity, and the decrease in SCD activity limits the ability of cancer cells within the tumor from adapting to a lower lipid environment (Fig. 4i). It is likely that these diet-mediated metabolic changes can cooperate with known alterations in whole-body metabolic regulation to mediate the effects of diet on tumor growth. Various low carbohydrate diets, including CR, intermittent fasting, and short-term fasting, that affect tumor progression alter whole-body metabolic regulation, most notably insulin signaling^7, 8, 10–16^. Of note, cancer cells harboring oncogenic PI3K pathway mutations were previously shown to be resistant to the tumor growth inhibitory effects of CR, an effect largely attributed to these cancer cells maintaining constitutive insulin signaling^10, 66^. Interestingly, SCD is a transcriptional target of SREBP1, which is activated downstream of PI3K by mTORC1^67, 68^, and up-regulation of SCD could thus contribute to the mechanisms by which oncogenic PI3K pathway mutations confer resistance to CR.

While these data provide a metabolic basis for how CR can impact tumor growth, it does not necessarily imply that CR should be recommended as a dietary intervention for cancer patients. Food restriction may not be tolerated in all patients, and weight loss can limit treatment options. Some studies have suggested a short-term but more extreme food restriction, especially prior to treatment with chemotherapy, immunotherapy, or other targeted agents, with the goal of enhancing the anti-cancer effects of the therapy to allow patients to benefit from the diet without the negative effects associated with long-term exposure to that diet^12, 69, 70^. Indeed, several clinical trials have been initiated to explore these dietary strategies^71–74^. Whether changes to lipid availability and metabolism, as well as other metabolic alterations, contribute to the effects of these short-term dietary interventions is an area for further exploration. However, this study demonstrates how diet can alter physiological levels of metabolites that are available to tumors and influence cancer cell metabolism to affect growth. A better understanding of these dietary effects on tumor metabolism and progression may lead to orthogonal strategies to mimic the effects of a particular diet, as well as provide guidance for how to best incorporate dietary interventions into the care of patients with cancer.

## Methods

### Cell lines and culture

AL1376 pancreatic ductal adenocarcinoma cells were isolated from C57BL/6J *LSL-Kras(G12D)*;*Trp53^fl/fl^*;*Pdx1-Cre* mice as previously described^75^. LGSP lung adenocarcinoma cells were derived from the C57BL/6J *LSL-Kras(G12D)*;*Trp53^fl/fl^*;*Ad-Cre* mouse lung cancer model as previously described^35, 36^. HeLa, Panc1, and A549 cells were obtained from the American Type Culture Collection (ATCC). All cells were cultured in DMEM (Corning Life Sciences) without pyruvate supplemented with 10% heat inactivated dialyzed fetal bovine serum (VWR) unless otherwise specified. Cells were passaged for no more than 6 months and routinely assayed for mycoplasma contamination.

### Animal studies

All experiments conducted in this study were approved by the MIT Committee on Animal Care (IACUC). For subcutaneous tumor growth, a maximum tumor burden of 2 cm^3^ was permitted per IACUC protocol, and these limits were not exceeded. All mice in this study were housed with a 12 h light and 12 h dark cycle and co-housed with littermates with ad libitum access to water, unless otherwise stated. All experimental groups were age-matched, numbered, and assigned based on treatment, and experiments were conducted in a blinded manner. Data was collected from distinct animals, where n represents biologically independent samples. Statistical methods were not performed to pre-determine sample size.

For subcutaneous tumors, C57BL/6J mice (The Jackson Laboratory 000664) were injected with 1-2 × 10^5^ mouse AL1376 cells as previously described^75^. Cells were injected subcutaneously into both flanks in 100 µl of phosphate-buffered saline (PBS) per injection. All mice were administered the AIN-93G control diet (Envigo TD.94045) during tumor formation, and 9-12 days after cell injection when palpable tumors had formed, animals were randomly placed into different diet groups. Mice were weighed before the start of diet administration to ensure different cohorts had similar starting body weights, and body weights were also measured over the course of each experiment. Tumor volume was determined using (π /6)(*W* ^2^)(*L*), where *W* represents width and *L* represents length as measured by calipers. At the end of each experiment, animals were euthanized, blood was collected by cardiac puncture, and tumors were rapidly harvested, weighed, and freeze-clamped in liquid nitrogen.

### Animal diets

The AIN-93G diet (Envigo TD.94045) was used as the control diet, and 40% caloric restriction (CR) and the ketogenic diet (KD) were formulated by modifying the AIN-93G diet as shown in Extended Data Table 1. CR (Envigo TD.170111) was formulated such that animals fed at 40% restriction consumed the same amounts of protein, fat, vitamins, and minerals, and only calories from carbohydrates were reduced by 40%. For CR studies, mice were individually housed. Prior to diet administration, the average daily consumption (by weight) of the AIN-93G control diet was determined. Upon experimental diet initiation, control mice were fed daily with the determined average daily food consumption weight, and calorie restricted mice were fed daily at 40% of the control food weight. The KD (Envigo TD.170112) was formulated by eliminating carbohydrates, reducing the protein component, and increasing fat content through the addition of lard and soybean oil. For KD studies, both control and KD groups were fed ad libitum. Average daily food consumption by weight and calories was measured over three days.

### Inhibitors

A939572 (Tocris 4845), GSK2194069 (Tocris 5303), and PF-05175157 (Tocris 5790) were used at the indicated concentrations.

### Plasmids and generation of stable cDNA expressing cells

For overexpression of SCD with a C-terminal HA tag, a lentivirus vector pLV[Exp]-Neo-CMV>hSCD/HA was constructed and ordered from VectorBuilder. tet-pLKO-Neo was used as an empty vector control. Stable cDNA expressing cell lines were generated by lentivirus infection for 24 h, followed by selection in DMEM containing 800 µg/ml G418. After selection, cells were maintained in 400 µg/ml G418 until used in experimental assays.

### Blood glucose and plasma insulin measurements

Blood glucose levels were measured using a Contour glucose meter (Ascensia Diabetes Care). Blood glucose from CR studies was assayed ∼20-24 hours after the previous meal. Blood glucose from KD studies was assayed at 8:00 am. Plasma insulin for CR and KD studies was sampled after a 4 h fast and was measured with an ultra-sensitive mouse insulin ELISA (Crystal Chem #90080).

### Tumor, plasma, and tumor interstitial fluid metabolite extraction

For tumors, frozen tissues were ground into powder using a mortar and pestle. Tissue powder was then weighed into glass vials (Thermofisher C4010-1, C4010-60BLK). Blood collected from animals was immediately placed in EDTA tubes (Sarstedt 41.1395.105) and centrifuged to separate plasma. Tumor interstitial fluid (TIF) was harvested as previously described^3^. For absolute quantification of polar metabolites in biofluids, plasma or TIF was mixed 1:1 with a solution of isotopically labeled amino acids, pyruvate, lactate, and β (Cambridge Isotope Laboratories, MSK-A2-1.2, CNLM-3819-H, CLM-1822-H, CLM-2440, CLM-10768, CLM-3853) in glass vials. Metabolites were extracted in 1.5 ml dichloromethane:methanol (containing 25 mg/L butylated hydroxytoluene, Sigma Aldrich B1378):0.88% KCl (w/v) (8:4:3), vortexed for 15 min, and centrifuged at maximum speed for 10 min. The extraction buffer contained 0.75 µg/ml norvaline and either 0.7 µg/ml tridecanoic acid or 0.7 µg/ml cis-10-heptadecenoic acid as internal standards. Polar metabolites (aqueous fraction) were transferred to Eppendorf tubes, dried under nitrogen gas, and stored at −80°C until further analysis. Lipids (organic fraction) were transferred to glass vials, dried under nitrogen gas, and immediately processed for analysis.

For glucose measurements in TIF, TIF was mixed 1:1 with a solution of isotopically labeled glucose at a known concentration (Cambridge Isotope Laboratories, CLM-1396). Samples were extracted in 300 µl of ice cold HPLC grade methanol, vortexed for 10 min, and centrifuged at maximum speed for 10 min. 280 µl of each extract was removed, dried under nitrogen gas, and stored at −80°C until further analysis.

### Cell culture isotopic labeling experiments and metabolite extraction

Cell lines were seeded at an initial density of 50,000-120,000 cells/well in a six-well dish in 2 mL of DMEM medium. Cells were incubated for 24 h and then washed three times with 2 mL of PBS. Cells were then incubated for 24 or 48 h in the indicated media and drug conditions. For isotope labeling experiments, cells were cultured in the presence of 10 mM U-^13^C_6_-glucose (Cambridge Isotope Laboratories, CLM-1396) and 1 mM U-^13^C_5_-glutamine (Cambridge Isotope Laboratories, CLM-1822-H) for 24 h.

Following the indicated treatments, media was aspirated from cells, and cells were rapidly washed in ice cold saline three times. The saline was aspirated, and 700 µl of methanol (containing 25 mg/L butylated hydroxytoluene, Sigma Aldrich B1378):0.88% KCl (w/v) (4:3) was added. Cells were scraped on ice, the extract was transferred to glass vials (Thermofisher C4010-1, C4010-60BLK), and 800 µl of dichloromethane was added. The final extraction buffer contained 0.75 µg/ml norvaline and either 0.7 µg/ml tridecanoic acid or 0.7 µg/ml cis-10-heptadecenoic acid as internal standards. The resulting extracts were vortexed for 15 min and centrifuged at maximum speed for 10 min. Lipids (organic fraction) were transferred to glass vials, dried under nitrogen gas, and immediately processed for analysis.

### Gas chromatography-mass spectrometry (GC-MS) analysis of polar metabolites

Polar metabolites were analyzed by GC-MS as described previously^76^. Dried and frozen metabolite extracts were derivatized with 16 µl of MOX reagent (Thermo Fisher TS-45950) for 60 min at 37°C, followed by derivatization with 20 µl of N-tert-butyldimethylsilyl-N-methyltrifluoroacetamide with 1% tert-butyldimethylchlorosilane (Sigma-Aldrich 375934) for 30 min at 60°C. Derivatized samples were analyzed by GC-MS, using a DB-35MS column (Agilent Technologies 122-3832) installed in an Agilent 7890B gas chromatograph coupled to an Agilent 5997B mass spectrometer. Helium was used as the carrier gas at a constant flow rate of 1.2 mL/min. One microliter of sample was injected in split mode (1:10) at 270°C. After injection, the GC oven was held at 100°C for 1 min, increased to 105°C at 2.5°C/min, held at 105°C for 2 min, increased to 250°C at 3.5°C/min, and then ramped to 320°C at 20°C/min. The MS system operated under electron impact ionization at 70 eV, and the MS source and quadrupole were held at 230°C and 150°C, respectively. The detector was used in scanning mode with an ion range of 100-650 m/z. Total ion counts were determined by integrating appropriate ion fragments for each metabolite^76^ using El-Maven software (Elucidata). Mass isotopologue distributions were corrected for natural abundance using IsoCorrectoR^77^. Metabolite data was normalized to the internal standard and biofluid volumes/tissue weights. Absolute concentrations of metabolites were calculated based on the known concentrations of isotopically labeled internal standards. Glucose in tumor interstitial fluid was analyzed by GC-MS as described previously^78^.

Dried and frozen metabolite extracts were derivatized with 50 µl of 2% (w/v) hydroxylamine hydrochloride in pyridine (Sigma-Aldrich) for 60 min at 90°C, followed by derivatization with 100 µl of propionic anhydride (Sigma-Aldrich) for 30 min at 60°C. Derivatized samples were then dried under nitrogen gas and resuspended in 100 µl of ethyl acetate (Sigma-Aldrich) in glass GC-MS vials. Samples were analyzed by GC-MS as described above, except helium was used as the carrier gas at a constant flow rate of 1.1 mL/min, and one microliter of sample was injected in splitless mode at 250°C. After injection, the GC oven was held at 80°C for 1 min, ramped to 280°C at 20°C/min, and held at 280°C for 4 min.

### Gas chromatography-mass spectrometry (GC-MS) analysis of fatty acid methyl esters

Fatty acid methyl esters (FAMEs) were analyzed by GC-MS. Dried lipid extracts were resuspended in 100 µl of toluene in glass vials and derivatized with 200 µl of 2% sulfuric acid in methanol overnight at 50°C. After derivatization, 500 µl of 5% NaCl was added, and FAMEs were extracted twice with 500 µl of hexane. Samples from cultured cell lines were dried under nitrogen gas, resuspended in 50 µl of hexane, and analyzed by GC-MS. Samples from animal tissues or biofluids were cleaned up with Bond Elut LRC-Florisil columns (Agilent Technologies 12113049). Columns were pre-conditioned with 3 mL of hexane, and then the FAME extracts in hexane were added to the column. FAMEs were finally eluted twice with 1 ml of hexane:diethyl ether (95:5 v/v), dried under nitrogen gas, and resuspended in hexane for GC-MS analysis. GC-MS was conducted with a DB-FastFAME column (Agilent Technologies G3903-63011) installed in an Agilent 7890A gas chromatograph coupled to an Agilent 5975C mass spectrometer. Helium was used as the carrier gas at a constant pressure of 14 psi. One microliter of sample was injected in splitless mode at 250°C. After injection, the GC oven was held at 50°C for 0.5 min, increased to 194°C at 25°C/min, held at 194°C for 1 min, increased to 245°C at 5°C/min, and held at 245°C for 3 min. The MS system operated under electron impact ionization at 70 eV, and the MS source and quadrupole were held at 230°C and 150°C, respectively. The detector was used in scanning mode with an ion range of 104-412 m/z. Total ion counts were determined by integrating appropriate ion fragments for each FAME using El-Maven software (Elucidata). Metabolite data was background corrected using a blank sample and normalized to the internal standard and biofluid volumes/tissue weights. Mass isotopologue distributions were corrected for natural abundance using IsoCorrectoR^77^. For calculation of relative fatty acid synthesis rates, the mass isotopologue distributions and total pool sizes of detected fatty acids were used in the fatty acid source analysis (FASA) script as previously described^37^. Raw data for fatty acid synthesis rate calculations are shown in Supplementary Table 1.

### Proliferation assays

Cells were seeded at an initial density of 20,000-40,000 cells/well in a 24-well dish in 1 mL of DMEM medium. Cells were incubated for 24 h, and cell number was assayed using sulforhodamine B staining, as previous described^79^, to establish the number of cells at the start of the experiment. Cells were then washed three times with 0.5 mL of PBS and incubated for 72 h in 1 mL of the indicated media and drug conditions. For evaluating the effects of changing media glucose concentrations on cell proliferation, media was changed daily to avoid depletion of glucose in the media. The number of cells at the end of the experiment was assayed using sulforhodamine B staining. Proliferation rate was determined using the following formula: Proliferation rate (doublings/day) = [Log2(Final Day 3 cell number/Initial Day 0 cell number)]/3 days.

### Preparation of de-lipidated fetal bovine serum and bovine serum albumin-fatty acid (BSA-FA) conjugates

Fetal bovine serum was stripped of lipids as previously described to generate de-lipidated cell culture media^34^. Exogenous fatty acids were supplemented in de-lipidated media as BSA-FA conjugates. Stock solutions of 0.17 mM BSA (0.17 mM), 0.17 mM BSA/1 mM 16:0, 0.17 mM BSA/1 mM 18:1(n-9), and 0.17 mM BSA/1 mM 18:2(n-6) were prepared as described by the Seahorse Bioscience protocol for the XF Analyzer. Solutions were filter-sterilized prior to addition to cell culture media at the indicated concentrations.

### Immunoblotting

Powdered tumor tissue was lysed in radioimmunoprecipitation assay (RIPA) buffer (Boston BioProducts BP-115-250) supplemented with Halt Protease and Phosphatase Inhibitor Cocktail (Thermo Fisher 78442) for 15 min at 4°C. Cell extracts were pre-cleared by centrifugation at maximum sipped for 10 min at 4°C, and protein concentration was measured with the Pierce BCA Protein Assay Kit (Thermo Scientific). For detection of SCD, lysates were not boiled. Lysates were resolved on SDS-PAGE and transferred electrophoretically to 0.2 µm nitrocellulose membranes (Bio-Rad) at 100 V for 60 min. The blots were blocked in Tris-buffered saline buffer (TBST; 10 mmol/L Tris-HCl, pH 8, 150 mmol/L NaCl, and 0.2% Tween-20) containing 5% (w/v) nonfat dry milk for 1 h, and then incubated with the specific primary antibody diluted in blocking buffer at 4°C overnight. Membranes were washed three times in TBST and incubated with HRP-conjugated secondary antibody for 1 h at room temperature. Membranes were washed three times and developed using enhanced chemiluminescence (Perkin-Elmer). Antibodies were used as follows: mouse Scd1 (Cell Signaling Technology #2438, 1:1000), human SCD (Abcam ab19862, 1:1000), HA-tag (Cell Signaling Technology #3724, 1:1000), and Vinculin (Cell Signaling Technology 13901, 1:1000). The mouse Scd1 antibody (Cell Signaling Technology #2438) was validated to preferentially recognize mouse Scd1 over human SCD, whereas the human SCD antibody (Abcam ab19862) was validated to only recognize human SCD.

### Statistics and reproducibility

Sample sizes, reproducibility, and statistical tests used for each figure are denoted in the figure legends. All graphs were generated using Graphpad Prism.

## Acknowledgments

We thank members of the Vander Heiden laboratory for useful discussions and experimental advice. E.C.L. was supported by the Damon Runyon Cancer Research Foundation (DRG-2299-17). Z.L. and K.M.S. were supported by NIH training grant T32GM007287. M.G.V.H. acknowledges support from the Emerald Foundation, the Lustgarten Foundation, a Faculty Scholar grant from the Howard Hughes Medical Institute, Stand Up To Cancer, the MIT Center for Precision Cancer Medicine, the Ludwig Center at MIT, and the NIH (R35CA242379, R01CA168653, R01CA201276, P30CA14051).

## Author contributions

Conceptualization, E.C.L. and M.G.V.H.; Methodology, E.C.L., Z.L., and M.G.V.H.; Investigation, E.C.L., A.M.W., and K.M.S.; Writing-Original Draft, E.C.L. and M.G.V.H.; Writing-Review and Editing, all authors; Supervision, E.C.L. and M.G.V.H.; Visualization, E.C.L.; Funding Acquisition, E.C.L. and M.G.V.H.

## Materials and Correspondence

Supplementary Information is available for this paper. Correspondence and requests for materials should be addressed to Matthew G. Vander Heiden.

---

**Extended Data Figure 1.**
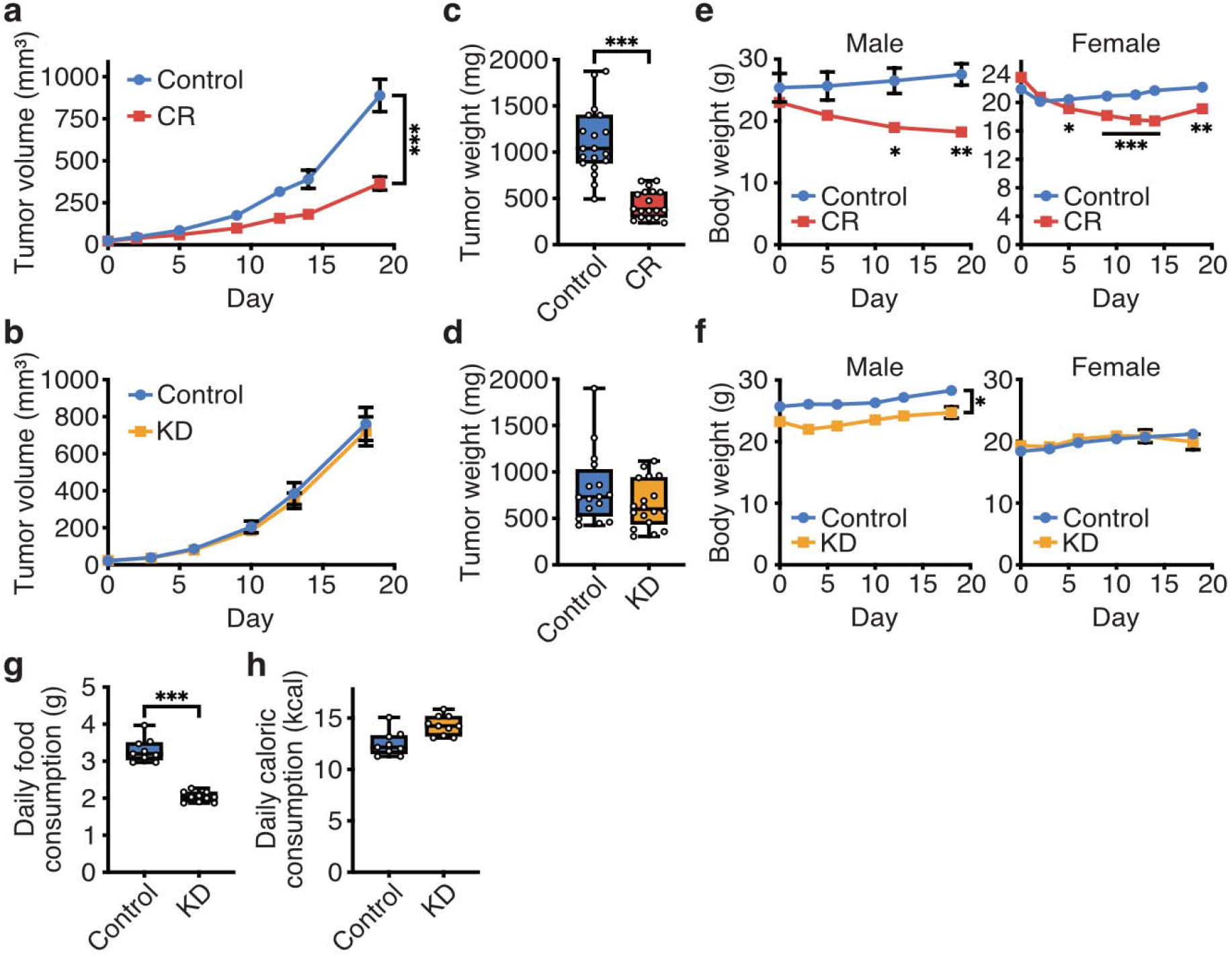
Caloric restriction, but not the ketogenic diet, impairs growth of pancreatic cancer allografts. **a**, **b**, Measured tumor volumes of subcutaneous AL1376 PDAC allografts in C57BL/6J mice exposed to (**a**) CR or a control diet, or (**b**) the KD or a control diet. Data are presented as mean ± SEM; (**a**) Control n = 4 male + 8 female mice, CR n = 3 male + 8 female mice; (**b**) Control n = 5 male + 4 female mice, KD n = 5 male + 5 female mice. Two-way repeated measures analysis of variance (ANOVA) was used for comparison between groups. **c**, **d**, Measured tumor weights of subcutaneous AL1376 allografts harvested at endpoint for the experiments shown in (**a-b**) where C57BL/6J mice were exposed to (**c**) CR or a control diet, or (**d**) the KD or a control diet. Data are presented as box-and-whisker plots displaying median and interquartile ranges; (**c**) Control n = 4 male + 8 female mice, CR n = 3 male + 8 female mice; (**d**) Control n = 5 male + 4 female mice, KD n = 5 male + 5 female mice. A two-tailed Student’s t-test was used for comparison between groups. **e**, **f**, Body weights of male and female C57BL/6J mice exposed to (**e**) CR or a control diet, or (**f**) the KD or a control diet. Data are presented as mean ± SEM; (**e**) Control n = 4 male + 8 female mice, CR n = 3 male + 8 female mice; (**f**) Control n = 5 male + 4 female mice, KD n = 5 male + 5 female mice. A two-tailed Student’s t-test was used for comparison between groups. **g**, **h**, Daily food consumption by weight (**g**) and daily caloric consumption (**h**) of the control diet and KD by C57BL/6J mice, as indicated. Data are presented as box-and-whisker plots displaying median and interquartile ranges; Control n = 5 male + 4 female mice, KD n = 5 male + 5 female mice. A two-tailed Student’s t-test was used for comparison between groups. *P < 0.05, **P < 0.01, ***P < 0.001.

**Extended Data Figure 2.**
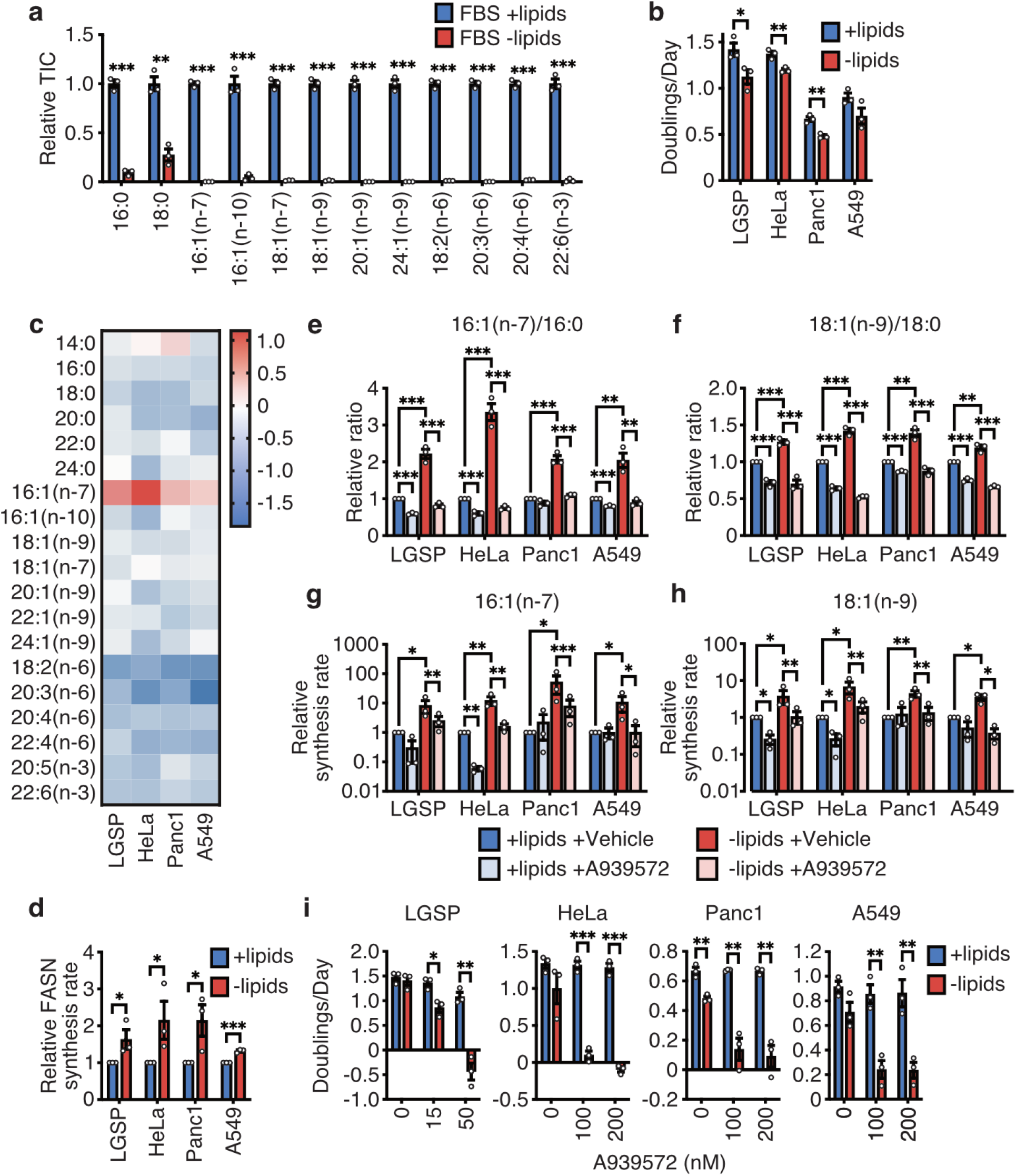
Increased stearoyl-CoA desaturase activity is required for cancer cells to adapt to exogenous lipid limitation. **a**, Levels of the indicated fatty acids in lipidated and de-lipidated fetal bovine serum (FBS) as specified. **b**, Doubling times of the specified cells cultured in media containing lipidated versus de-lipidated serum. **c**, Fold changes in levels of the indicated fatty acids extracted from cells cultured in de-lipidated versus lipidated media for 24 h. **d**, Cells were cultured in media containing U-^13^C-glucose and U-^13^C-glutamine for 24 h in lipidated versus de-lipidated media as indicated, and ^13^C incorporation into fatty acids was detected by GC-MS (see Extended Data Fig. 3). The mass isotopologue distributions and total pool sizes of detected fatty acids were used to calculate a FASN synthesis rate by fatty acid source analysis (FASA, see Supplementary Table 1). A one-tailed Student’s t-test was used for comparison between groups. **e**, **f**, Ratios of 16:1(n-7)/16:0 (**e**) and 18:1(n-9)/18:0 (**f**) fatty acids extracted from cells cultured in lipidated versus de-lipidated media with or without the SCD inhibitor A939572 (LGSP: 50 nM; HeLa, Panc1, A549: 200 nM) for 24 h as indicated. **g**, **h**, 16:1(n-7) (**g**) and 18:1(n-9) (**h**) synthesis rates, calculated as described in (**c**) (see also Supplementary Table 1), in cells cultured in lipidated versus de-lipidated media with or without A939572 (LGSP: 50 nM; HeLa, Panc1, A549: 200 nM) for 24 h as indicated. A one-tailed ratio paired t-test was used for comparison between groups. **i**, Doubling times of the specified cells cultured in lipidated versus de-lipidated media with the indicated concentrations of A939572. All data are presented as mean ± SEM; unless otherwise indicated, n = 3 biologically independent replicates, and a two-tailed Student’s t-test was used for comparison between groups unless otherwise noted above. *P < 0.05, **P < 0.01, ***P < 0.001.

**Extended Data Figure 3.**
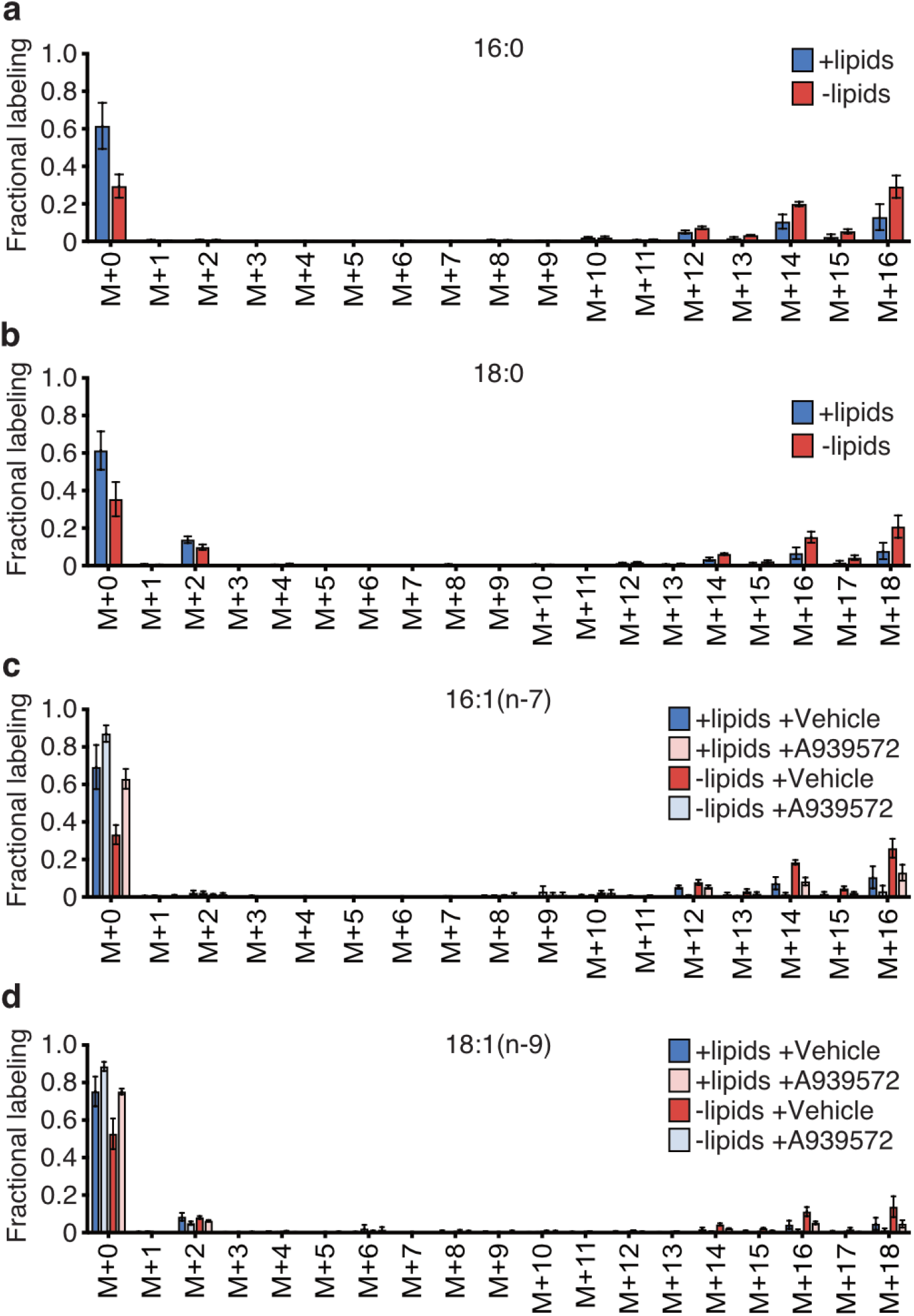
Representative mass isotopologue distributions of fatty acids labeled by U-^13^C-glucose and U-^13^C-glutamine. **a**, **b**, Mass isotopologue distributions of 16:0 (**a**) and 18:0 (**b**) from AL1376 cells labeled with U-^13^C-glucose and U-^13^C-glutamine for 24 h in lipidated versus de-lipidated media. **c**, **d**, Mass isotopologue distributions of 16:1(n-7) (**c**) and 18:1(n-9) (**d**) from AL1376 cells labeled with U-^13^C-glucose and U-^13^C-glutamine for 24 h in lipidated versus de-lipidated media containing 50 nM A939572. All data are presented as mean ± SEM; n = 3 biologically independent replicates. These data were used to calculate fatty acid synthesis rates in Fig. 3, Extended Data Fig. 2, Extended Data Fig. 5, and Extended Data Fig. 6; see also Supplementary Table 1 for raw data.

**Extended Data Figure 4.**
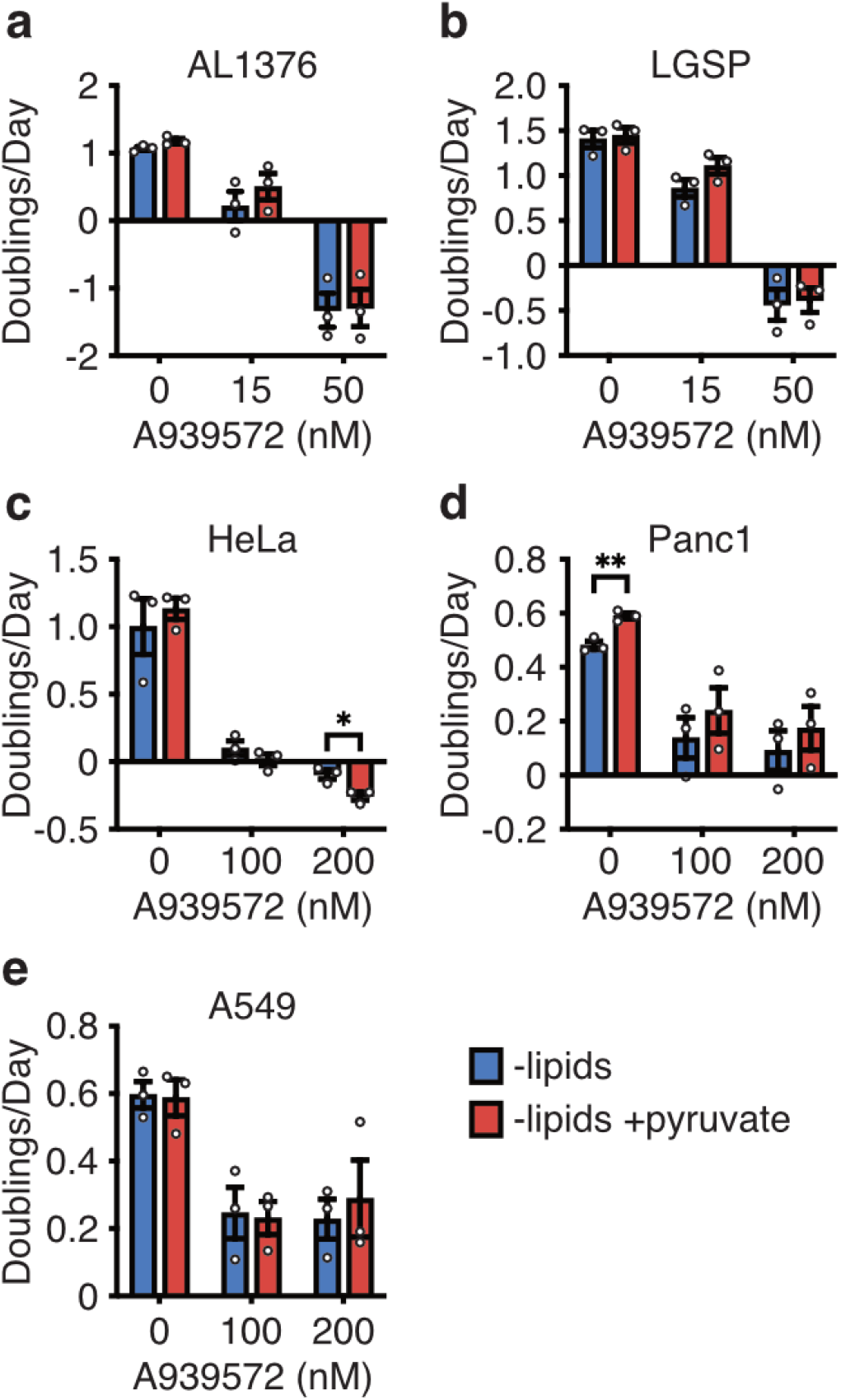
Stearoyl-CoA desaturase inhibition does not impair cell proliferation via NAD+ limitation. **a-e**, Doubling times of the indicated cell lines cultured in de-lipidated media containing the indicated concentrations of the SCD inhibitor A939572 and 1 mM pyruvate. All data are presented as mean ± SEM; n = 3 biologically independent replicates. A two-tailed Student’s t-test was used for comparison between groups. *P < 0.05, **P < 0.01, ***P < 0.001.

**Extended Data Figure 5.**
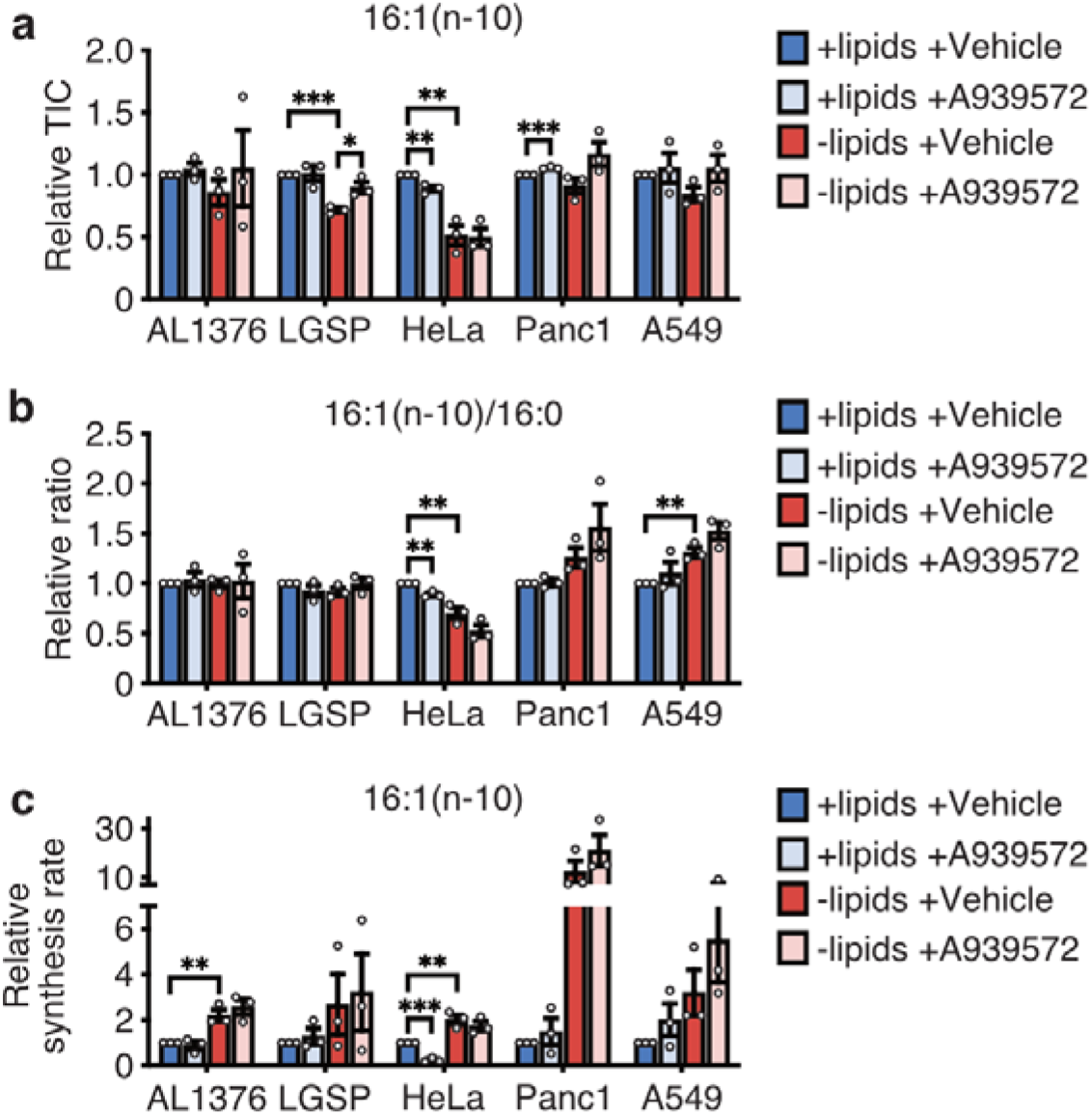
Stearoyl-CoA desaturase inhibition does not increase production of 16:1(n-10) in these cells. **a**, **b**, 16:1(n-10) (**a**) and 16:1(n-10)/16:0 (**b**) levels in extracts from cells cultured in lipidated versus de-lipidated media, with or without the SCD inhibitor A939572 (AL1376, LGSP: 50 nM; HeLa, Panc1, A549: 200 nM) for 24 h. **c**, Cells were cultured in the presence of U-^13^C-glucose and U-^13^C-glutamine for 24 h in lipidated versus de-lipidated media, with or without A939572 (AL1376, LGSP: 50 nM; HeLa, Panc1, A549: 200 nM), and ^13^C incorporation into 16:1(n-10) was assessed by GC-MS. The mass isotopologue distribution and total pool size of 16:1(n-10) were used to calculate a 16:1(n-10) synthesis rate by fatty acid source analysis (FASA, see Supplementary Table 1). All data are presented as mean ± SEM; n = 3 biologically independent replicates. A two-tailed Student’s t-test was used for comparison between groups. *P < 0.05, **P < 0.01, ***P < 0.001.

**Extended Data Figure 6.**
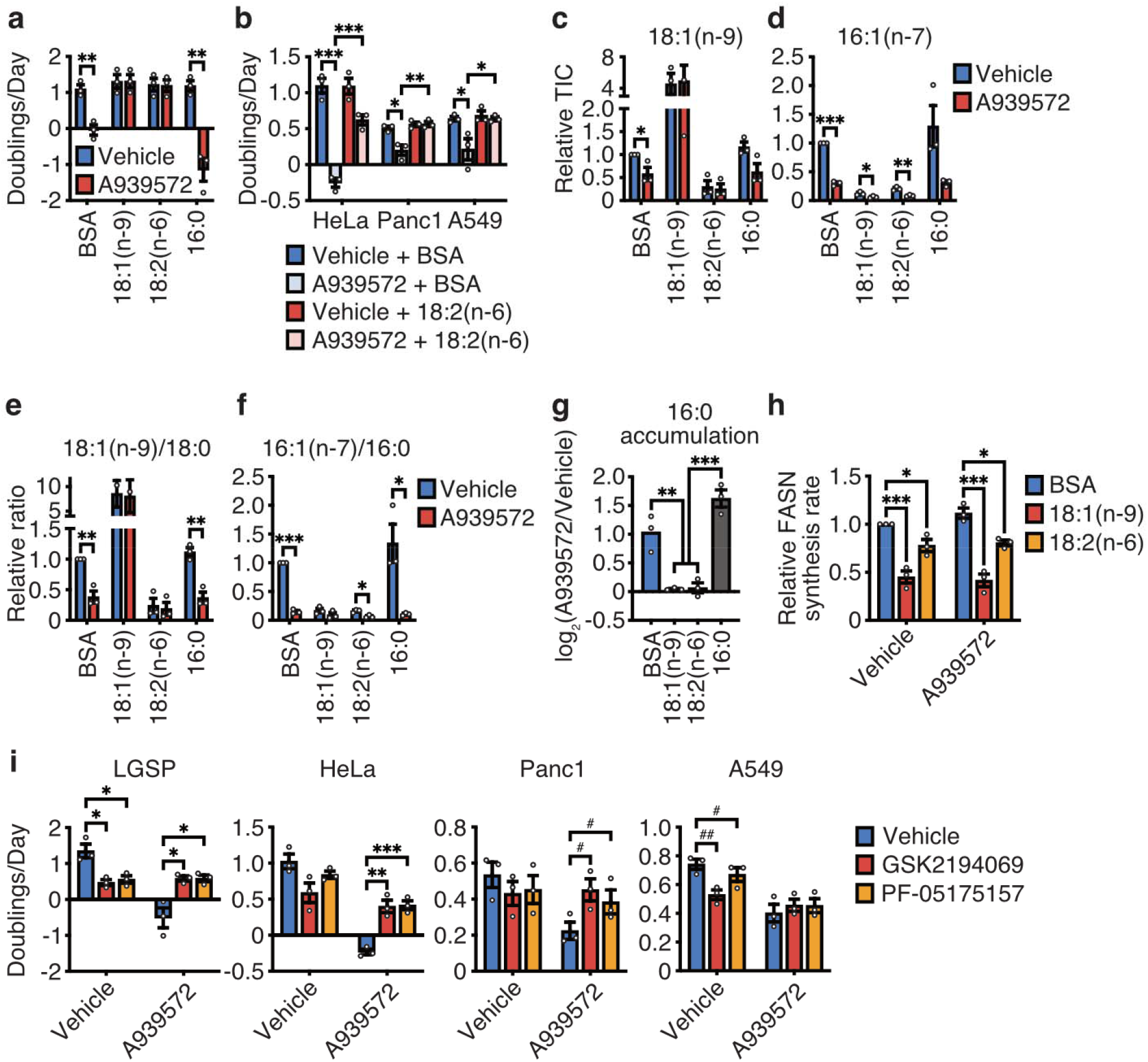
Stearoyl-CoA desaturase inhibition impairs cell proliferation by inducing palmitate accumulation. **a**, Doubling times of LGSP cells cultured in de-lipidated media with or without 50 nM of the SCD inhibitor A939572, BSA, 100 µM 18:1(n-9), 100 µM 18:2(n-6), or 15 µM 16:0 as indicated. **b**, Doubling times of cells cultured in de-lipidated media containing 200 nM A939572, BSA, or 25 µM 18:2(n-6). **c-f**, 18:1(n-9) (**c**), 16:1(n-7) (**d**), 18:1(n-9)/18:0 (**e**), and 16:1(n-7)/16:0 (**f**) levels in extracts from LGSP cells cultured in de-lipidated media with or without 50 nM A939572, BSA, 100 µM 18:1(n-9), 100 µM 18:2(n-6), or 15 µM 16:0 for 48 h as indicated. **g**, Fold change in levels of 16:0 induced by exposure to 50 nM A939572 in LGSP cells cultured in de-lipidated media containing BSA, 100 µM 18:1(n-9), 100 µM 18:2(n-6), or 15 µM 16:0 for 48 h as indicated. **h**, LGSP cells were cultured in the presence of U-^13^C-glucose and U-^13^C-glutamine for 24 h in de-lipidated media containing 50 nM A939572, BSA, 100 µM 18:1(n-9), or 100 µM 18:2(n-6), and ^13^C incorporation into fatty acids was detected by GC-MS. The mass isotopologue distributions and total pool sizes of detected fatty acids were used to calculate a FASN synthesis rate by fatty acid source analysis (FASA, see Supplementary Table 1). **i**, Doubling times of cells cultured in de-lipidated media containing A939572 (LGSP: 50 nM; HeLa, Panc1, A549: 200 nM), the FASN inhibitor GSK2194069 (LGSP: 0.3 µM; HeLa, A549: 0.015 µM; Panc1: 0.05 µM), or the ACC inhibitor PF-05175157 (LGSP: 10 µM; HeLa: 0.3 µM; Panc1: 1 µM; A549: 0.5 µM) as specified. All data are presented as mean ± SEM; n = 3 biologically independent replicates. *P < 0.05, **P < 0.01, ***P < 0.001 by a two-tailed Student’s t-test. ^#^P < 0.05, ^##^P < 0.01 by a paired two-tailed Student’s t-test.

**Extended Data Figure 7.**
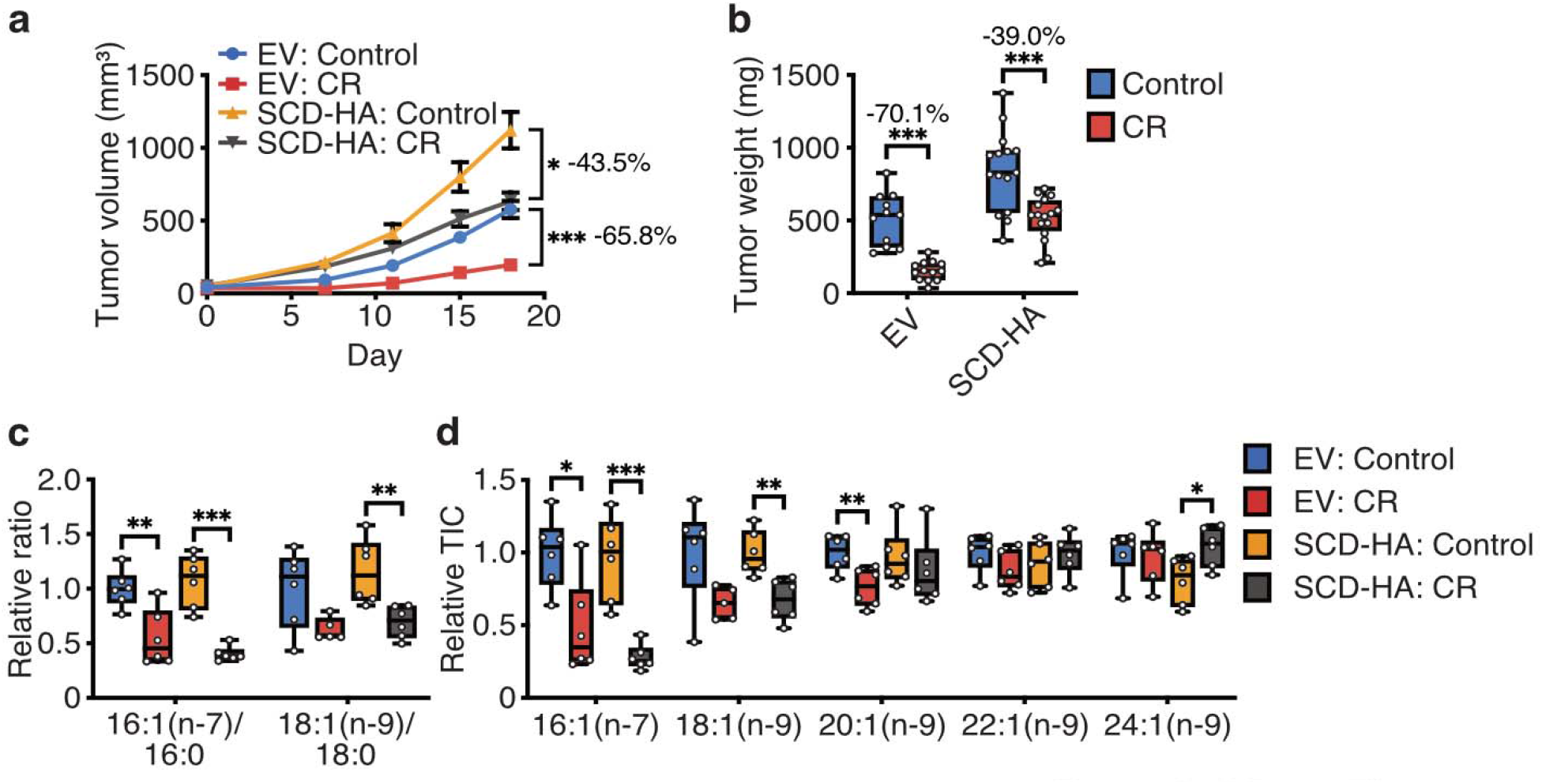
Caloric restriction inhibits tumor growth by limiting exogenous lipid availability and impairing tumor SCD activity. **a**, **b**, Measured tumor volumes (**a**) and tumor weights (**b**) of subcutaneous AL1376 allograft tumors expressing empty vector (EV) or SCD-HA in C57BL/6J mice administered a control or CR diet as indicated. Data are presented as (**a**) mean ± SEM and (**b**) box-and-whisker plots displaying median and interquartile ranges; EV: Control n = 3 male + 3 female mice, EV: CR n = 4 male + 4 female mice, SCD-HA: Control n = 4 male + 4 female mice, SCD-HA: CR n = 4 male + 4 female mice. Two-way repeated measures analysis of variance (ANOVA) was used for comparison between groups. **c**, **d**, MUFA/SFA ratios (**c**) and MUFA levels (**d**) in subcutaneous AL1376 allograft tumors expressing EV or SCD-HA harvested from C57BL/6J mice administered a control or CR diet for 18 days as indicated. Data are presented as box-and-whisker plots displaying median and interquartile ranges; n = 6 per group. A two-tailed Student’s t-test was used for comparison between groups. *P < 0.05, **P < 0.01, ***P < 0.001.

**Extended Data Table 1.**
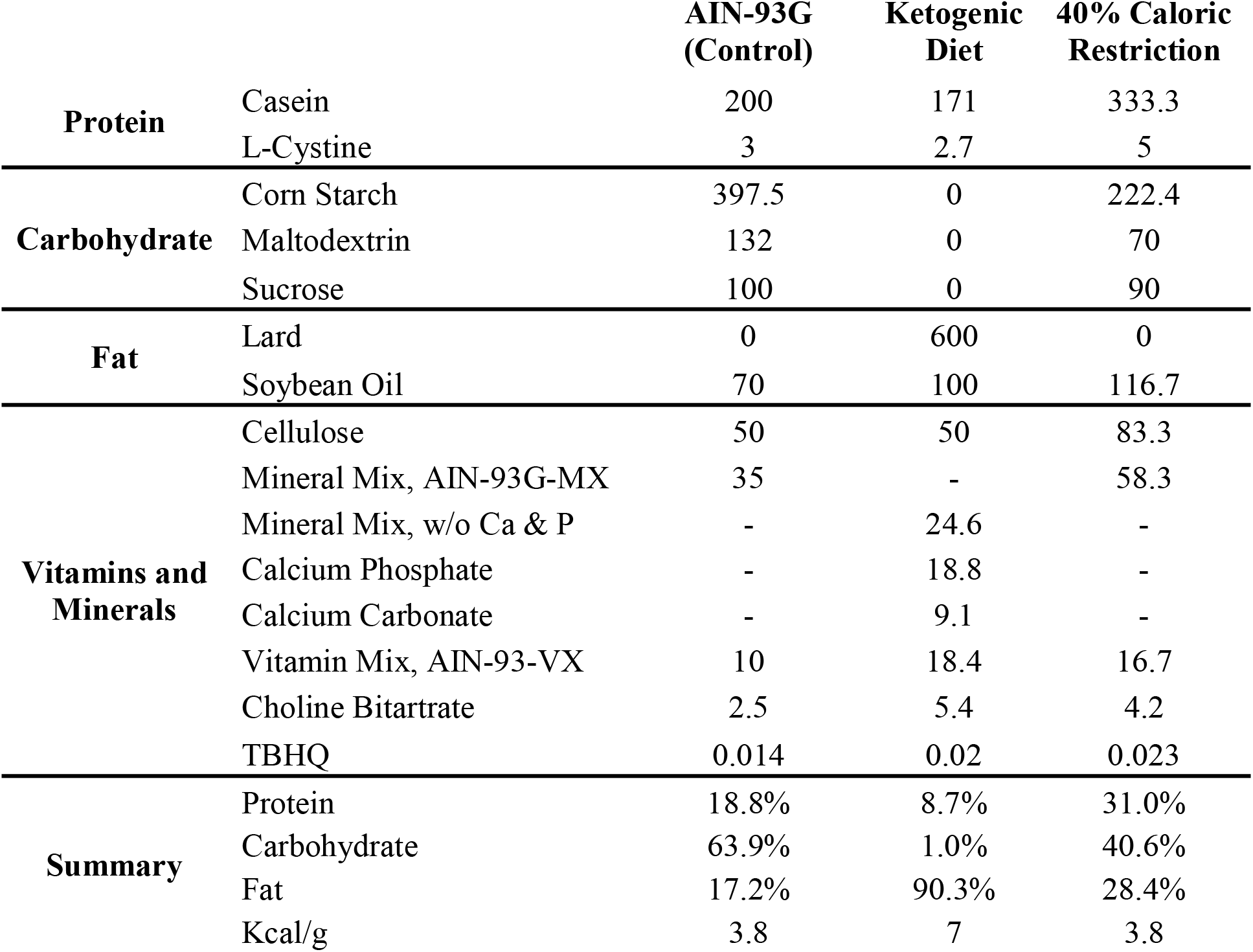
Compositions of diets used in this study.

**Supplementary Table 1. Raw data used for fatty acid source analysis (FASA) synthesis rate calculations in Fig. 3, Extended Data Fig. 2, Extended Data Fig. 5, and Extended Data Fig. 6.**

[see attached .xlsx file]

## References

1. Lien, E. C. & Vander Heiden, M. G. A framework for examining how diet impacts tumour metabolism. Nat Rev Cancer (2019).

2. Sullivan, M. R. et al. Increased Serine Synthesis Provides an Advantage for Tumors Arising in Tissues Where Serine Levels Are Limiting. Cell Metab 29, 1410–1421.e4 (2019).

3. Sullivan, M. R. et al. Quantification of microenvironmental metabolites in murine cancers reveals determinants of tumor nutrient availability. Elife 8, (2019).

4. Maddocks, O. D. et al. Serine starvation induces stress and p53-dependent metabolic remodelling in cancer cells. Nature 493, 542–546 (2013).

5. Maddocks, O. D. K. et al. Modulating the therapeutic response of tumours to dietary serine and glycine starvation. Nature 544, 372–376 (2017).

6. Gao, X. et al. Dietary methionine influences therapy in mouse cancer models and alters human metabolism. Nature (2019).

7. Meynet, O. & Ricci, J.-E. Caloric restriction and cancer: molecular mechanisms and clinical implications. Trends Mol Med 20, 419–427 (2014).

8. Lee, C. & Longo, V. D. Fasting vs dietary restriction in cellular protection and cancer treatment: from model organisms to patients. Oncogene 30, 3305–3316 (2011).

9. Poff, A. M., Ari, C., Seyfried, T. N. & D’Agostino, D. P. The ketogenic diet and hyperbaric oxygen therapy prolong survival in mice with systemic metastatic cancer. PLoS One 8, e65522 (2013).

10. Kalaany, N. Y. & Sabatini, D. M. Tumours with PI3K activation are resistant to dietary restriction. Nature 458, 725–731 (2009).

11. Hopkins, B. D. et al. Suppression of insulin feedback enhances the efficacy of PI3K inhibitors. Nature (2018).

12. Nencioni, A., Caffa, I., Cortellino, S. & Longo, V. D. Fasting and cancer: molecular mechanisms and clinical application. Nat Rev Cancer 18, 707–719 (2018).

13. Nogueira, L. M., Lavigne, J. A., Chandramouli, G. V., Lui, H., Barrett, J. C. & Hursting, S. D. Dose-dependent effects of calorie restriction on gene expression, metabolism, and tumor progression are partially mediated by insulin-like growth factor-1. Cancer Med 1, 275–288 (2012).

14. Lu, Z. et al. Fasting selectively blocks development of acute lymphoblastic leukemia via leptin-receptor upregulation. Nat Med 23, 79–90 (2017).

15. Beyaz, S. et al. High-fat diet enhances stemness and tumorigenicity of intestinal progenitors. Nature 531, 53–58 (2016).

16. Hursting, S. D., Dunlap, S. M., Ford, N. A., Hursting, M. J. & Lashinger, L. M. Calorie restriction and cancer prevention: a mechanistic perspective. Cancer Metab 1, 10 (2013).

17. Muir, A., Danai, L. V., Gui, D. Y., Waingarten, C. Y., Lewis, C. A. & Vander Heiden, M. G. Environmental cystine drives glutamine anaplerosis and sensitizes cancer cells to glutaminase inhibition. Elife 6, e27713 (2017).

18. Cantor, J. R. et al. Physiologic Medium Rewires Cellular Metabolism and Reveals Uric Acid as an Endogenous Inhibitor of UMP Synthase. Cell 169, 258–272.e17 (2017).

19. Tardito, S. et al. Glutamine synthetase activity fuels nucleotide biosynthesis and supports growth of glutamine-restricted glioblastoma. Nat Cell Biol 17, 1556–1568 (2015).

20. Vande Voorde, J., et al. Improving the metabolic fidelity of cancer models with a physiological cell culture medium. Sci Adv 5, eaau7314 (2019).

21. Muir, A. & Vander Heiden, M. G. The nutrient environment affects therapy. Science 360, 962–963 (2018).

22. Warburg, O. On the origin of cancer cells. Science 123, 309–314 (1956).

23. Vander Heiden, M. G. & DeBerardinis, R. J. Understanding the Intersections between Metabolism and Cancer Biology. Cell 168, 657–669 (2017).

24. Ben-Haim, S. & Ell, P. 18F-FDG PET and PET/CT in the evaluation of cancer treatment response. J Nucl Med 50, 88–99 (2009).

25. Bardeesy, N. et al. Both p16(Ink4a) and the p19(Arf)-p53 pathway constrain progression of pancreatic adenocarcinoma in the mouse. Proc Natl Acad Sci U S A 103, 5947–5952 (2006).

26. Danai, L. V. et al. Altered exocrine function can drive adipose wasting in early pancreatic cancer. Nature 558, 600–604 (2018).

27. Douris, N. et al. Adaptive changes in amino acid metabolism permit normal longevity in mice consuming a low-carbohydrate ketogenic diet. Biochim Biophys Acta 1852, 2056–2065 (2015).

28. Zhang, J. et al. Low ketolytic enzyme levels in tumors predict ketogenic diet responses in cancer cell lines in vitro and in vivo. J Lipid Res 59, 625–634 (2018).

29. Collet, T. H. et al. A Metabolomic Signature of Acute Caloric Restriction. J Clin Endocrinol Metab 102, 4486–4495 (2017).

30. Miller, K. N. et al. Aging and caloric restriction impact adipose tissue, adiponectin, and circulating lipids. Aging Cell 16, 497–507 (2017).

31. Raeini-Sarjaz, M., Vanstone, C. A., Papamandjaris, A. A., Wykes, L. J. & Jones, P. J. Comparison of the effect of dietary fat restriction with that of energy restriction on human lipid metabolism. Am J Clin Nutr 73, 262–267 (2001).

32. Kulkarni, S. R., Armstrong, L. E. & Slitt, A. L. Caloric restriction-mediated induction of lipid metabolism gene expression in liver is enhanced by Keap1-knockdown. Pharm Res 30, 2221–2231 (2013).

33. Yu, D. et al. Calorie-Restriction-Induced Insulin Sensitivity Is Mediated by Adipose mTORC2 and Not Required for Lifespan Extension. Cell Rep 29, 236–248.e3 (2019).

34. Hosios, A. M., Li, Z., Lien, E. C. & Vander Heiden, M. G. Preparation of Lipid-Stripped Serum for the Study of Lipid Metabolism in Cell Culture. Bio-protocol 8, e2876 (2018).

35. DuPage, M., Dooley, A. L. & Jacks, T. Conditional mouse lung cancer models using adenoviral or lentiviral delivery of Cre recombinase. Nat Protoc 4, 1064–1072 (2009).

36. Gocheva, V. et al. Quantitative proteomics identify Tenascin-C as a promoter of lung cancer progression and contributor to a signature prognostic of patient survival. Proc Natl Acad Sci U S A 114, E5625–E5634 (2017).

37. Argus, J. P. et al. Development and Application of FASA, a Model for Quantifying Fatty Acid Metabolism Using Stable Isotope Labeling. Cell Rep 25, 2919–2934.e8 (2018).

38. Peter, A. et al. Hepatic lipid composition and stearoyl-coenzyme A desaturase 1 mRNA expression can be estimated from plasma VLDL fatty acid ratios. Clin Chem 55, 2113–2120 (2009).

39. Sjögren, P. et al. Fatty acid desaturases in human adipose tissue: relationships between gene expression, desaturation indexes and insulin resistance. Diabetologia 51, 328–335 (2008).

40. Kim, W. et al. Polyunsaturated Fatty Acid Desaturation Is a Mechanism for Glycolytic NAD^+^ Recycling. Cell Metab (2019).

41. Sullivan, L. B., Gui, D. Y., Hosios, A. M., Bush, L. N., Freinkman, E. & Vander Heiden, M. G. Supporting Aspartate Biosynthesis Is an Essential Function of Respiration in Proliferating Cells. Cell 162, 552–563 (2015).

42. Birsoy, K., Wang, T., Chen, W. W., Freinkman, E., Abu-Remaileh, M. & Sabatini, D. M. An Essential Role of the Mitochondrial Electron Transport Chain in Cell Proliferation Is to Enable Aspartate Synthesis. Cell 162, 540–551 (2015).

43. Titov, D. V., Cracan, V., Goodman, R. P., Peng, J., Grabarek, Z. & Mootha, V. K. Complementation of mitochondrial electron transport chain by manipulation of the NAD+/NADH ratio. Science 352, 231–235 (2016).

44. Diehl, F. F., Lewis, C. A., Fiske, B. P. & Vander Heiden, M. G. Cellular redox state constrains serine synthesis and nucleotide production to impact cell proliferation. Nat Metab 1, 861–867 (2019).

45. Gui, D. Y. et al. Environment Dictates Dependence on Mitochondrial Complex I for NAD+ and Aspartate Production and Determines Cancer Cell Sensitivity to Metformin. Cell Metab 24, 716–727 (2016).

46. Roongta, U. V. et al. Cancer cell dependence on unsaturated fatty acids implicates stearoyl-CoA desaturase as a target for cancer therapy. Mol Cancer Res 9, 1551–1561 (2011).

47. Kamphorst, J. J. et al. Hypoxic and Ras-transformed cells support growth by scavenging unsaturated fatty acids from lysophospholipids. Proc Natl Acad Sci U S A 110, 8882–8887 (2013).

48. Peck, B. et al. Inhibition of fatty acid desaturation is detrimental to cancer cell survival in metabolically compromised environments. Cancer Metab 4, 6 (2016).

49. Mason, P. et al. SCD1 inhibition causes cancer cell death by depleting mono-unsaturated fatty acids. PLoS One 7, e33823 (2012).

50. Vriens, K. et al. Evidence for an alternative fatty acid desaturation pathway increasing cancer plasticity. Nature 566, 403–406 (2019).

51. Ariyama, H., Kono, N., Matsuda, S., Inoue, T. & Arai, H. Decrease in membrane phospholipid unsaturation induces unfolded protein response. J Biol Chem 285, 22027–22035 (2010).

52. Paumen, M. B., Ishida, Y., Muramatsu, M., Yamamoto, M. & Honjo, T. Inhibition of carnitine palmitoyltransferase I augments sphingolipid synthesis and palmitate-induced apoptosis. J Biol Chem 272, 3324–3329 (1997).

53. Piccolis, M. et al. Probing the Global Cellular Responses to Lipotoxicity Caused by Saturated Fatty Acids. Mol Cell 74, 32–44.e8 (2019).

54. Zhu, X. G. et al. CHP1 Regulates Compartmentalized Glycerolipid Synthesis by Activating GPAT4. Mol Cell 74, 45–58.e7 (2019).

55. DeBose-Boyd, R. A. & Ye, J. SREBPs in Lipid Metabolism, Insulin Signaling, and Beyond. Trends Biochem Sci 43, 358–368 (2018).

56. Hardwicke, M. A. et al. A human fatty acid synthase inhibitor binds β the keto-substrate site. Nat Chem Biol 10, 774–779 (2014).

57. Griffith, D. A. et al. Decreasing the rate of metabolic ketone reduction in the discovery of a clinical acetyl-CoA carboxylase inhibitor for the treatment of diabetes. J Med Chem 57, 10512–10526 (2014).

58. Ventura, R. et al. Inhibition of de novo Palmitate Synthesis by Fatty Acid Synthase Induces Apoptosis in Tumor Cells by Remodeling Cell Membranes, Inhibiting Signaling Pathways, and Reprogramming Gene Expression. EBioMedicine 2, 808–824 (2015).

59. Heuer, T. S. et al. FASN Inhibition and Taxane Treatment Combine to Enhance Anti-tumor Efficacy in Diverse Xenograft Tumor Models through Disruption of Tubulin Palmitoylation and Microtubule Organization and FASN Inhibition-Mediated Effects on Oncogenic Signaling and Gene Expression. EBioMedicine 16, 51–62 (2017).

60. Svensson, R. U. et al. Inhibition of acetyl-CoA carboxylase suppresses fatty acid synthesis and tumor growth of non-small-cell lung cancer in preclinical models. Nat Med 22, 1108–1119 (2016).

61. Peck, B. & Schulze, A. Lipid desaturation - the next step in targeting lipogenesis in cancer. FEBS J 283, 2767–2778 (2016).

62. Röhrig, F. & Schulze, A. The multifaceted roles of fatty acid synthesis in cancer. Nat Rev Cancer 16, 732–749 (2016).

63. Pinkham, K. et al. Stearoyl CoA Desaturase Is Essential for Regulation of Endoplasmic Reticulum Homeostasis and Tumor Growth in Glioblastoma Cancer Stem Cells. Stem Cell Reports 12, 712–727 (2019).

64. Young, R. M. et al. Dysregulated mTORC1 renders cells critically dependent on desaturated lipids for survival under tumor-like stress. Genes Dev 27, 1115–1131 (2013).

65. Ackerman, D. et al. Triglycerides Promote Lipid Homeostasis during Hypoxic Stress by Balancing Fatty Acid Saturation. Cell Rep 24, 2596–2605.e5 (2018).

66. Curry, N. L. et al. Pten-null tumors cohabiting the same lung display differential AKT activation and sensitivity to dietary restriction. Cancer Discov 3, 908–921 (2013).

67. Düvel, K. et al. Activation of a metabolic gene regulatory network downstream of mTOR complex 1. Mol Cell 39, 171–183 (2010).

68. Peterson, T. R. et al. mTOR complex 1 regulates lipin 1 localization to control the SREBP pathway. Cell 146, 408–420 (2011).

69. Raffaghello, L. et al. Starvation-dependent differential stress resistance protects normal but not cancer cells against high-dose chemotherapy. Proc Natl Acad Sci U S A 105, 8215–8220 (2008).

70. Lee, C. et al. Fasting cycles retard growth of tumors and sensitize a range of cancer cell types to chemotherapy. Sci Transl Med 4, 124ra27 (2012).

71. de Groot, S. et al. The effects of short-term fasting on tolerance to (neo) adjuvant chemotherapy in HER2-negative breast cancer patients: a randomized pilot study. BMC Cancer 15, 652 (2015).

72. Dorff, T. B. et al. Safety and feasibility of fasting in combination with platinum-based chemotherapy. BMC Cancer 16, 360 (2016).

73. Safdie, F. M. et al. Fasting and cancer treatment in humans: A case series report. Aging (Albany NY) 1, 988–1007 (2009).

74. Bauersfeld, S. P. et al. The effects of short-term fasting on quality of life and tolerance to chemotherapy in patients with breast and ovarian cancer: a randomized cross-over pilot study. BMC Cancer 18, 476 (2018).

75. Mayers, J. R. et al. Tissue of origin dictates branched-chain amino acid metabolism in mutant Kras-driven cancers. Science 353, 1161–1165 (2016).

76. Lewis, C. A. et al. Tracing compartmentalized NADPH metabolism in the cytosol and mitochondria of mammalian cells. Mol Cell 55, 253–263 (2014).

77. Heinrich, P. et al. Correcting for natural isotope abundance and tracer impurity in MS-, MS/MS- and high-resolution-multiple-tracer-data from stable isotope labeling experiments with IsoCorrectoR. Sci Rep 8, 17910 (2018).

78. Antoniewicz, M. R., Kelleher, J. K. & Stephanopoulos, G. Measuring deuterium enrichment of glucose hydrogen atoms by gas chromatography/mass spectrometry. Anal Chem 83, 3211–3216 (2011).

79. Vichai, V. & Kirtikara, K. Sulforhodamine B colorimetric assay for cytotoxicity screening. Nat Protoc 1, 1112–1116 (2006).

